# Count your bits: fingerprint benchmarking to assess broad chemical space representation

**DOI:** 10.1101/2025.06.16.659994

**Authors:** Florian Huber, Julian Pollmann

## Abstract

Quantifying molecular similarity is a cornerstone of cheminformatics, underpinning applications from virtual screening and nearest-neighbor search to chemical space visualization and the evaluation of machine-learning predictions. Although Tanimoto-comparisons of 2D fingerprints are widely used, the practical behavior of these similarity measures depends strongly on the fingerprint type, representation (binary vs. count), and whether fingerprints are folded into fixed-length vectors. Here, we systematically benchmark a broad set of common fingerprint types, including dictionary-based, circular (Morgan/FCFP), path-based (RDKit), topological-distance-based (Atom Pair), hybrid distance-encoded (MAP4), torsion, LINGO, and Avalon, across multiple large datasets and complementary evaluation tasks. We quantify fingerprint specificity via duplicate rates and mass discrepancies, characterize score distributions and compound-size dependence, assess top-k ranking agreement, and compare fingerprint similarities to a graph-based reference. Across benchmarks, count (and often log-count) variants generally improve specificity and structural alignment, while folding-induced bit collisions can strongly distort similarities for high-occupancy fingerprints, making unfolded variants particularly important for RDKit and often necessary for MAP4 on heterogeneous datasets. To support reproducible benchmarking and future extensions, we introduce the open-source Python library chemap, providing unified computation of folded, unfolded, and frequency-folded fingerprints and optimized similarity calculations.

**Scientific contribution:** We introduce a multi-criteria benchmarking framework for molecular fingerprints that goes beyond classical virtual-screening retrieval tests by evaluating specificity (fingerprint duplicates), score behavior (including compound-size dependence), ranking agreement, and neighborhood structure on large, heterogeneous small-molecule datasets. Our results reveal that common default settings, especially folding for high-occupancy fingerprints, can introduce severe bit-collision artifacts, and that count (often log-count) and unfolded variants substantially improve specificity and agreement with structure-based references. We release the open-source Python library chemap to standardize these fingerprint variants and enable reproducible, extensible benchmarking for future fingerprint development.

## Introduction

Quantitative measures of molecular similarity (or distance) are essential for a wide range of applications in cheminformatics. They are used to identify structurally related molecules in large databases, to rank candidate compounds in virtual screening workflows ^1,2^ and to perform high-level data analysis tasks such as dimensionality reduction and chemical space visualization ^3–6^. Molecular similarity scores are further increasingly used as a target for training and evaluating deep-learning models for the prediction of molecular structures from mass spectrometry or NMR data ^7–9^.

At the heart of molecular similarity is the question: *What does it mean for two molecules to be similar?* Unfortunately, there is no single, universally valid definition of molecular similarity, but rather a multitude of molecular similarity measures, each tailored to specific data analysis goals or property predictions^10,11^. Two molecules may be considered similar if they share synthetic pathways, exhibit closely related biological or physicochemical properties, or differ only slightly in their overall molecular structure. Determining similarity thus heavily depends on the context and intended application.

For example, researchers working on lipidomics may focus more on broad structural classes, while paying less attention to minor variations in fatty acid chains. In contrast, others may be interested in the exact length and saturation levels of side chains, adopting a viewpoint in which even small structural alterations can be significant.

Given these complexities, it is unsurprising that a variety of computational methods, fingerprint algorithms, and similarity metrics have been proposed ^12–17^. The prevailing approach generally involves first computing a molecular fingerprint as an abstract representation of a compound before applying a similarity or distance metric to these fingerprints. Among the numerous available metrics, the Tanimoto coefficient (closely related to the Jaccard index for binary data), Dice, and Cosine similarities are commonly employed ^2,18^. Tanimoto scores have become a de facto standard, frequently cited as an effective choice for fingerprint-based similarity comparisons ^1,2,13,18^.

Commonly applied fingerprint algorithms, such as those based on circular or path-based methods, come with several favorable aspects. They can typically be computed very efficiently. Once generated, comparing large numbers of compounds by Tanimoto similarity is also highly scalable. Moreover, unlike data-driven embeddings derived from neural networks or other machine learning models, these classical fingerprints are model-independent. The bit patterns they produce retain a level of interpretability since certain bits can often be traced back to defined structural fragments in the molecule.

Nonetheless, Tanimoto-based similarities also have well-known limitations ^6,19^. Even subtle structural changes can sometimes lead to unexpectedly large shifts in similarity values.

Conversely, changes can be scored as irrelevant even though a chemist would classify them as relevant. Furthermore, most fingerprint algorithms may collapse distinct molecules into identical or near-identical bit patterns, resulting in no effective discrimination.

These shortcomings are not inherent to the Tanimoto metric alone, but rather, they stem from the interaction between the chosen metric and the underlying fingerprint design. The common practice of broadly referring to all such measures as “Tanimoto similarity” can mask the important variability introduced by different fingerprint representations and parameter choices ^12^. Moreover, the choice of the fingerprint algorithm as well as the similarity metric is often perceived as arbitrary ^20^.

One reason for this might be that a well-informed decision on the “right” fingerprint for a specific task is often difficult to make. There is a large body of research on fingerprint types and variants going back many decades. Systematic fingerprint evaluation approaches are dominated by a focus on retrieval tasks such as the search for molecules of the same activity class within a larger set of non-active compounds or decoys ^1,12,21^. This, however, only highlights one perspective on molecular representations. Yet, molecular fingerprints are also foundational for many other tasks such as chemical space visualizations ^5,17^, even on datasets with a very diverse chemical composition ^6^, and as input (or target) for a quickly growing number of machine learning models ^8,22–26^. For those tasks, extensive and systematic benchmarking approaches are largely missing. Finally, the aspect of larger datasets in combination with advanced computational capabilities induces important shifts with respect to technical limitations, but also underlying statistical effects, making it worthwhile to re-evaluate “traditional” fingerprint choices and settings.

The objective of this study is to provide a systematic analysis of common fingerprint types and their adjustable variants with respect to a broad range of tasks on large-scale datasets of small molecules. This is accompanied by a new Python library, chemap, as a basis for expansion by the community and adoption for comparing and benchmarking future fingerprint algorithms.

### Different molecular fingerprint classes

In this work, the focus is placed on 2D fingerprints, where structural features are encoded as high-dimensional binary or count vectors. Over the last decades a large variety of different fingerprints with near endless possible variants has been proposed ^13,17,20,27^. Among the most commonly used fingerprint categories are **dictionary-based** fingerprints which encode molecules along a set of predefined substructures. Common examples, which will also be used here, are MACCS keys ^28^, PubChem fingerprints ^29^, Klekota-Roth fingerprints ^30^, and Biosynfoni^16^.

Instead of using predefined sets of substructures, many other fingerprints identify (or count) substructures based on defined criteria. The following distinction can be made between **local substructure-based** methods, most importantly: circular fingerprints, which encode local neighborhoods of atoms within a specified radius ^2,27,31^ (we here use Morgan and FCFP fingerprints), and path-based fingerprints, which capture fragments defined by linear paths up to a certain length ^2^, here using the RDKit fingerprint ^32^. A widely used special case is the topological torsion fingerprint, which hash fixed-length atom sequences (typically four atoms) based on their atom-type pattern ^33^. A related, string-inspired alternative is given by LINGO fingerprints, which represent molecules via overlapping SMILES substrings ^34^.

Beyond purely local fragments, **topological-distance-based** approaches such as Atom-Pair fingerprints ^35^ describe pairs of atoms together with their graph distance.A more recent method, the MAP4 fingerprint ^14^, can be seen as a hybrid between local-substructure and topological distance. Finally, Avalon fingerprints provide a widely used, practical hashed fingerprint family that aggregates a broad set of substructure and path-like features into a fixed-length bit vector^36^.

All these fingerprints can, in principle, be computed in various flavors. A binary representation will only determine presence or absence of a specific substructure, while count representations include also its number of occurrences in a given molecule. While dictionary-based fingerprints naturally come as a fixed-length vector, for the other fingerprints the number of identified unique substructures will depend on the fingerprint settings and the given data. Most commonly, those fingerprints are folded onto a fixed vector size. Depending on the implementation, however, it is also possible to omit the folding step to compute unfolded fingerprints, typically with each substructure being represented in a hashed form.

Yet another category of fingerprints are pharmacophore fingerprints, which we will ignore in this work due to their much larger computational costs (for 3D variants) and their relatively poor performance on broader activity classification tasks ^17^.

An alternative to fragment-based representations is graph-based similarity measures, such as those derived from the Maximum Common Subgraph (MCS) or Maximum Common Edge Subgraph (MCES), which can offer a more complete structural comparison ^37^. Such measures are conceptually very appealing and capable of avoiding several pitfalls of common molecular fingerprints, which is why we will use them in this work as a point of reference. Yet these measures might also fail to reflect the nuanced notion of similarity that a chemist might apply. By focusing primarily on the largest common subgraph, these methods emphasize shared structure over subtle but potentially crucial differences. For a pair of molecules sharing an extensive scaffold, for instance, replacing a methyl group with a strongly electron-withdrawing nitro substituent can dramatically change properties such as reactivity or binding affinity, yet the MCS-based similarity score would remain high.

In practice, however, the much more relevant limitation is the very high computational cost of such graph-based metrics. Although it is possible to improve overall computation times using threshold or approximated approaches ^6,37^, these remain many orders of magnitude slower than fingerprint-based similarity computations. The need to solve an NP-hard task results in poor scaling of such algorithms for larger and more complex molecules ^38^, thereby rendering them unsuitable for large-scale similarity calculations for comparing larger molecules.

Recently, a different class of data-driven approaches has emerged, leveraging deep learning and other machine learning techniques to learn abstract representations of molecules, e.g., embeddings ^39–41^. Such embeddings can capture both global and local structural features in a data-driven manner. These approaches typically require large, well-annotated datasets and may lack the interpretability found in classical fingerprints. But they offer a novel and promising route to compare, analyze and visualize large chemical datasets ^42^. In this work, however, we will solely focus on model-independent similarity computations.

### Evaluation of molecular similarity measures

Evaluating different molecular similarity measures in order to guide the selection of appropriate methods and parameters is non-trivial ^17^. A frequent use case of molecular fingerprints has long been the search for compounds of similar activity ^43^ which is commonly evaluated by the retrieval of compounds of the same activity class ^1,12,15,21^ or by their ability to serve as a basis for activity classification models ^17,39,44^. At times, the high variability in datasets and fingerprint types led to inclusive results ^45^ making it difficult to decide in favor of or against a particular type of fingerprint, and recent efforts began to systematically re-evaluate quantitative structure-activity relationships (QSAR) modeling on natural product datasets ^17^. Those approaches reveal important aspects regarding QSAR, but in the end this is only one possible perspective to assess molecular representations. However, such benchmarking represents only one particular task, which, in our opinion, does not reflect the full scale of current applications of molecular similarity. It is commonly applied for the task of detecting active compounds in larger datasets also containing decoys. While this might represent certain virtual screening tasks well enough, it does not necessarily reflect a universal requirement for good chemical similarity measures.

Additionally, focusing on the top-ranked candidates will emphasize precision in a narrow subset of all possible molecule pairs. This leaves broader aspects of similarity assessment underexplored, as methods that excel at identifying a few key hits may not perform equally well in more general contexts.

Besides screening tasks, however, there is a wide range of different applications for molecular fingerprints. They are frequently used to create chemical space visualizations ^4,5,17,46^ with the specific aim of revealing how well a measure preserves meaningful structural relationships across the entire space. However, quantitative comparison of dimensionality-reduction-based visualizations is inherently difficult and therefore often not ideal for systematic benchmarking of fingerprint types and variants

Molecular fingerprints are further used to predict chemical classes ^16,26^ or taxonomic origin ^47^. While NPClassifier uses Morgan count fingerprints without large-scale comparisons to other fingerprint types ^26^, Biosynfoni, which was mainly designed to capture biosynthetic distances, was also evaluated using a compound classification task ^16^ and includes benchmarking against few other fingerprints (MACCS, Morgan, RDKit). Those comparisons are not easily generalizable though, because they were limited to only seven chemical classes and further only included the most basic binary MACCS, Morgan, and RDKit fingerprints. Capecchi and Reymond ^47^ focused on the MAP4 fingerprint ^21^ but also included a small comparison to few other (binary) fingerprints. This, too, is not systematic enough to serve as a general benchmark but does add yet another possible evaluation approach.

Finally, there is an increasing use of molecular fingerprints in complex machine-learning tasks where such fingerprint representations are used as key model input ^22,24–26,44^, as prediction target ^25,48^, or to evaluate compound predictions or generations, for instance with respect to ground truth molecules ^8,49^. The latter use as a distance measure to a ground truth is conceptually very different from retrieval-based tasks. While retrieval focuses entirely on top-ranked candidates, thereby emphasizing precision in a narrow subset of all possible molecule pairs, the use as a general-purpose target or loss function also further requires producing meaningful numbers of dissimilarity.

We propose a benchmarking framework that evaluates fingerprint variants across (i) specificity/collisions, (ii) score behavior and size dependence, (iii) ranking agreement, and (iv) downstream predictive and neighborhood-structure tasks.

## Methods

The following methodology was devised for the purpose of benchmarking multiple fingerprints across a range of chemical contexts:

First several datasets were utilized and created. Secondly, molecular fingerprints were examined in detail, and a new benchmarking library, chemap, was developed. This library can be used to generate folded, unfolded and frequency-scaled fingerprints. Finally, we outline an evaluation framework for different fingerprint variations. This framework utilizes similarity metrics, bioactivity prediction and chemical space visualization.

### Datasets

Organic molecules inherently display a huge diversity with large variations in the estimates of how many molecules it contains ^3^. In this study, we use various datasets drawn from natural products and metabolomics research, each selected for specific benchmarking tasks.

#### ms2structures dataset (37,811 compounds)

We assembled a curated collection of compounds measured and annotated by tandem mass spectrometry. We merged the training and evaluation data from MS2Deepscore, a deep-learning model for predicting chemical similarity from mass spectra^50^ with the overlapping benchmarking set, MassSpecGym ^8^ to create a dataset of 37,811 unique compounds.

#### biostructures dataset (718,067 compounds)

We chose the biostructures dataset from Kretschmer et al. ^6^, a large, biologically relevant and chemically heterogeneous collection previously used for large-scale chemical space analysis, as a deliberate stress test for broad chemical-space representation (in contrast to screening-focused, predominantly drug-like collections such as ChEMBL^51^ or ZINC^52^, or very general repositories such as PubChem ^29^. We removed 30 entries from the 718,097 compounds in the dataset that RDKit could not convert to fingerprints. Using the Classyfire API ^53^ we added chemical class information for 695,152 compounds.

#### 25-subclasses dataset (75,000 compounds)

Of 25 of the 27 most common chemical subclasses, excluding Glycerophosphoglycerophosphoglycerols and Triradylcglycerols since they were too easily distinguishable from the rest, 3000 compounds were randomly sampled from the biostructures dataset. This results in a balanced classification dataset.

#### 120-subclasses dataset (120,000 compounds)

This is another classification task oriented dataset, now with a random sample of 1000 unique compounds for the 120 most common chemical subclasses from the biostructures dataset.

#### rascalMCES dataset (5,413,677 compound pairs)

We randomly sampled 5,557,963 compound pairs from the ms2structures dataset whose precursor masses differ by at most 100 Da. RascalMCES scores ^37^ were computed with RDKit ^32^ on an Intel Core i9-13900K (settings: similarityThreshold=0.05, maxBondMatchPairs=1000, minFragSize=3, timeout=60 s). After excluding 144,286 timed-out pairs, our final benchmarking set contained 5,413,677 pairs.

#### bioactivity dataset (2,680 compounds, 1,340 of which with one or more activities)

Taking data from Boldini et al ^17^, a new dataset was assembled. First, all compounds with at least one specified bioactivity (out of 12 possible classes) were selected from their dataset. Second, an equivalent quantity of compounds without any such activity was incorporated into the non-active cases. Compounds can have specified bioactivity multiple categories at the same time. In addition, the labels are highly unbalanced with the rarest class having 20 instances and the most frequent class having 364 instances.

All datasets are publicly available at https://doi.org/10.5281/zenodo.18682051.

### Molecular Fingerprints

Most molecular fingerprints are computed using chemap, which is a new Python library we created to compute fingerprints from RDKit ^32^, Scikit-Fingerprints ^54^. Chemap further contains own implementations of MAP4 as well as Lingo. The MAP4 implementation was created based on the original source code ^21^, the scikit-fingerprints implementation ^54^ and an implementation from https://github.com/LucaCappelletti94/map4. Lingo was implemented based on the scikit-fingerprints implementation. This was necessary, because the existing implementations did not provide all variant options at the same time (binary, count, folded and unfolded). Further, we used the Biosynfoni fingerprint ^16^.

All code used for fingerprint computations can be found on GitHub: https://github.com/florian-huber/molecular_fingerprint_comparisons

The chemap library is available at: https://github.com/matchms/chemap

### Folded, Unfolded, Frequency Folded, and Sparse fingerprints

In their default settings, molecular fingerprints are folded into fixed-sized numerical vectors. An exception is dictionary-based fingerprints, such as the here used MACCS, PubChem fingerprint, Klekota-Roth and Biosynfoni, which all have pre-defined sets of substructures that determine their vector size. For all other fingerprints, the default **folded** variant with a vector size of 4096 was used, unless otherwise indicated. Those vectors can be computed to be binary (presence/absence) or count vectors. In addition, we work with **unfolded** fingerprints. For those, the detected substructures are hashed to 32-bit integers, which will avoid virtually all bit collisions while still being computationally reasonably efficient. Fingerprints are in those cases either stored as array of substructure hashes (for binary fingerprints) or as tuples of substructure hashes and the corresponding count values.

Not to be confused with unfolded formats are “**sparse**” formats, by which we here mean specific datatypes provided by SciPy ^55^ such as csr arrays that efficiently store and handle sparse arrays without storing any zeros. Within the library chemap, we also provide functions to convert non-folded fingerprints into csr arrays.

Finally, we also tested **frequency folded** fingerprints by selecting the most frequently occurring unfolded fingerprint bits within a larger dataset. Unlike the typical folding procedure, e.g., by using modulo, this will avoid any bit collisions in the final vector. This, however, comes at the price of ignoring lower frequency bits.

### Scaling, and Bit-Weighing

Many fingerprints are rooted in the detection of unique subsets of a molecular graph. Each unique subset is thereby treated equally, which has the advantage that the algorithms do not need to incorporate extensive heuristics to account for chemical knowledge. However, this will also not distinguish very common, or trivial, chemical subsets from more particular subsets. This problem resembles a common challenge in Natural Language Processing, where simply counting the occurrences of every word in a sentence is often insufficient. This is because many words, such as so-called stopwords, carry little specific meaning, while others are key for determining the topic or mood of the sentence.

We here adapt the concept of term frequency (TF) as well as the inverse document frequency (IDF) to account for the overall relevance of individual fingerprint bits in a fingerprint. The term frequency is here identical to the values in a count fingerprint, which represent the number of occurrences of a specific substructure in a molecule. The IDF is computed in a normalized way as follows:

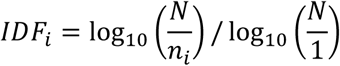

Where *N* is the total number of compounds in the dataset and *n*_*i*_ is the number of compounds that contain the substructure corresponding to bit *i*.

An additional, dataset-independent approach is the log-scaling of count values. This is applied to account for the fact that absolute differences between small numbers (count 1 to count 2, for instance) might sometimes better be considered as more relevant than the same shift in higher counts (e.g., count 20 to 21).

### Fingerprint Similarity Metrics

We applied Tanimoto scores to compute the similarity between molecular fingerprints. For binary fingerprints X and Y this meant using the following formula which is commonly referred to as Tanimoto or Jaccard index.

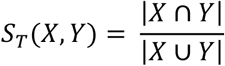

For count vectors X and Y containing positive integers as well as for scaled vectors containing float numbers, we use the closest relative which is often termed generalized Tanimoto coefficient or counted Tanimoto similarity (other terms are generalized Jaccard index or Ruzicka score):

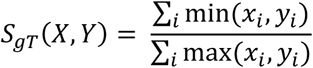

When used with binary fingerprints, the generalized Tanimoto coefficient leads to the exact same results as the original Tanimoto coefficient. In the following, we will therefore simply refer to “Tanimoto” without further distinction. For optimal runtime, however, we used different Tanimoto implementations for folded binary fingerprints, folded count fingerprints, unfolded binary fingerprints and unfolded count fingerprints. These functions were optimized by just-in-time compilation and parallelization using Numba^56^. All used Python code is available in the library “chemap”, with all source code on Github: https://github.com/matchms/chemap

### Percentile scores

To facilitate comparison between different fingerprint types and datasets, we convert raw similarity values into percentile scores on a 0–100 scale per fingerprint per dataset. Given a symmetric similarity matrix *S* ∈ ℝ^*N*×*N*^, we extract the strictly upper-triangular entries, rank them (using average ranks for ties), and then linearly scale the ranks so that the smallest similarity corresponds to 0 and the largest to 100. This transformation preserves the relative ordering of similarity scores while placing them on a uniform, interpretable scale. This makes cross-dataset and cross-fingerprint comparisons more meaningful.

### RascalMCES scores

To compare how well fingerprint-based similarities align with RascalMCES, we used Spearman correlation to assess how consistently the two methods order compound pairs, without being influenced by the actual score values or their distributions.

### Biological activity prediction models

For the bioactivity prediction task, fingerprints were evaluated in a multilabel setting where each compound is associated with a multi-hot vector of bioactivity labels (one binary label per target/property). For every fingerprint type and variant, SMILES strings were converted into fixed-length fingerprint vectors using the respective fingerprint generator. For selected fingerprints, an additional “collapse-free” variant was evaluated by generating unfolded fingerprints and subsequently reducing dimensionality to 4096 features by retaining the most frequently non-zero features across the dataset. Count-based fingerprints were transformed using a log(1+x) mapping, while binary fingerprints were used without transformation.

A simple feed-forward neural network, implemented using PyTorch ^57^, was trained to predict all labels simultaneously. The model consisted of a single hidden layer (3000 units) followed by batch normalization and dropout (p = 0.2). We intentionally used a single simple architecture to probe representational sufficiency rather than maximize predictive performance. Training used binary cross-entropy. To address label imbalance, per-label positive class weights were computed from the training data only and provided to the loss function.

Hyperparameters were selected separately for each fingerprint using a simple grid search over learning rate, batch size, and weight decay. To minimize chance effects during tuning and ensure comparability across fingerprints, a single fixed tuning split was generated once and reused for all fingerprint configurations. The tuning split was created by stratifying samples according to label density (number of active labels per compound, capped at three bins) and then partitioning into train and validation subsets; an additional holdout partition was retained but not used for the tuning objective. Each grid configuration was trained for up to 30 epochs with early stopping (patience 6). The primary tuning objective was macro-averaged ROC-AUC on the tuning validation set; if macro-AUC was undefined (e.g., due to missing positives/negatives for some labels), micro-averaged F1 served as a fallback objective.

After selecting the best hyperparameter combination per fingerprint, final performance was estimated using repeated stratified random splits (n = 5). For each split, the dataset was partitioned into train/validation/test subsets using the same label-density stratification strategy. Models were trained for up to 50 epochs with early stopping and then evaluated on the held-out test set. The code used for training and evaluation of the models can be found on Github: https://github.com/florian-huber/molecular_fingerprint_comparisons

### Chemical subclass prediction models

For different fingerprint types and variants, a simple neural network was trained on the task of predicting one out of 120 possible chemical subclasses. For each fingerprint configuration, molecules were converted into fixed-length fingerprint vectors using the respective fingerprint generator. In addition to standard folded fingerprints (fixed dimensional hashing), we also evaluated a “collision-free” variant for selected fingerprints: fingerprints were first generated in an unfolded representation and subsequently reduced to a fixed dimensionality by retaining the 4096 most frequently non-zero features across the dataset.

Subclass labels were encoded as integer class indices. Model performance was assessed using stratified k-fold cross-validation (k = 3) in order to reduce split-dependent variance. In each fold, the training portion was further split into a training and validation subset using stratified sampling to enable early stopping. Using PyTorch ^57^ a feed-forward multilayer perceptron (MLP) with four hidden layers (input ⃗ 4096 units ⃗ 2048 units ⃗ 2048 units ⃗ 1024 units ⃗ softmax over 120 subclasses) was trained using cross-entropy loss. Batch normalization and dropout (p = 0.2) were applied after the hidden layers. Optimization was performed with AdamW (learning rate 1×10⁻³, weight decay 1×10⁻⁴, batch size 512). All experiments were run with a fixed random seed for reproducibility. The code used for training and evaluation is available on Github: https://github.com/florian-huber/molecular_fingerprint_comparisons.

For comparison, we also trained a much simpler model only containing one hidden layer (3000 units), the results of which are displayed in the supplemental material.

### Subclass consistency evaluation

Subclass consistency was evaluated using the 25-subclasses dataset (3000 compounds for each of 25 subclasses, see above dataset section). Using Pynndescent ^58^ an approximate nearest neighbor graph (k=100) was computed for each fingerprint type and variant using chemap’s tanimoto distance implementation.

### Dimensionality reduction chemical space plots

For visualizations of chemical space, UMAP ^59^ was used as dimensionality reduction technique to compute 2D coordinates of all compounds. This was done based on the respective fingerprint type and variant. In chemap, we provide all code necessary to compute UMAP coordinates using the generalized Tanimoto distance, which is simply the inverse of the Tanimoto similarity, for either folded or unfolded fingerprint variants.

In addition, we provide a GPU-optimized faster variant that relies on cuml (https://github.com/rapidsai/cuml)^60^ in chemap. This, however, only supports folded fingerprints and cosine as a similarity measure.

We assessed the class separability of each UMAP embedding using a supervised “local structure” proxy adapted from Huang et al. ^61^. For each fingerprint variant, we trained a radial-basis-function support vector machine (SVM) on the 2D UMAP coordinates using the compound Subclass annotation as class label (25 classes total). We used a stratified train/test split with 80% of samples for training and 20% for testing, and repeated the evaluation for three different random seeds. Reported performance is the mean (and variability) of the held-out classification accuracy across the three runs.

## Results

To explore the practical impact of different fingerprints, we first compared their bit occupations across a dataset of 718,067 biomolecular structures (**biostructures dataset**). Figure 1 provides an overview of the average bit occupation for various fingerprint types. As expected, clear differences emerged. The RDKit path-based fingerprint, computed with a default sequence length of 7, tends to yield a high bit occupation rate, particularly in the smaller, 1024-bit vectors. Here, about 90 % of all fingerprint bits are occupied for more than 40% of all compounds in the biomolecular dataset. By contrast, Morgan or FCFP fingerprints exhibit significantly lower bit occupancy. Even for small 1024-bit fingerprints, only about 2% of all bits are occupied for more than 40% of all compounds.

**Figure 1.**
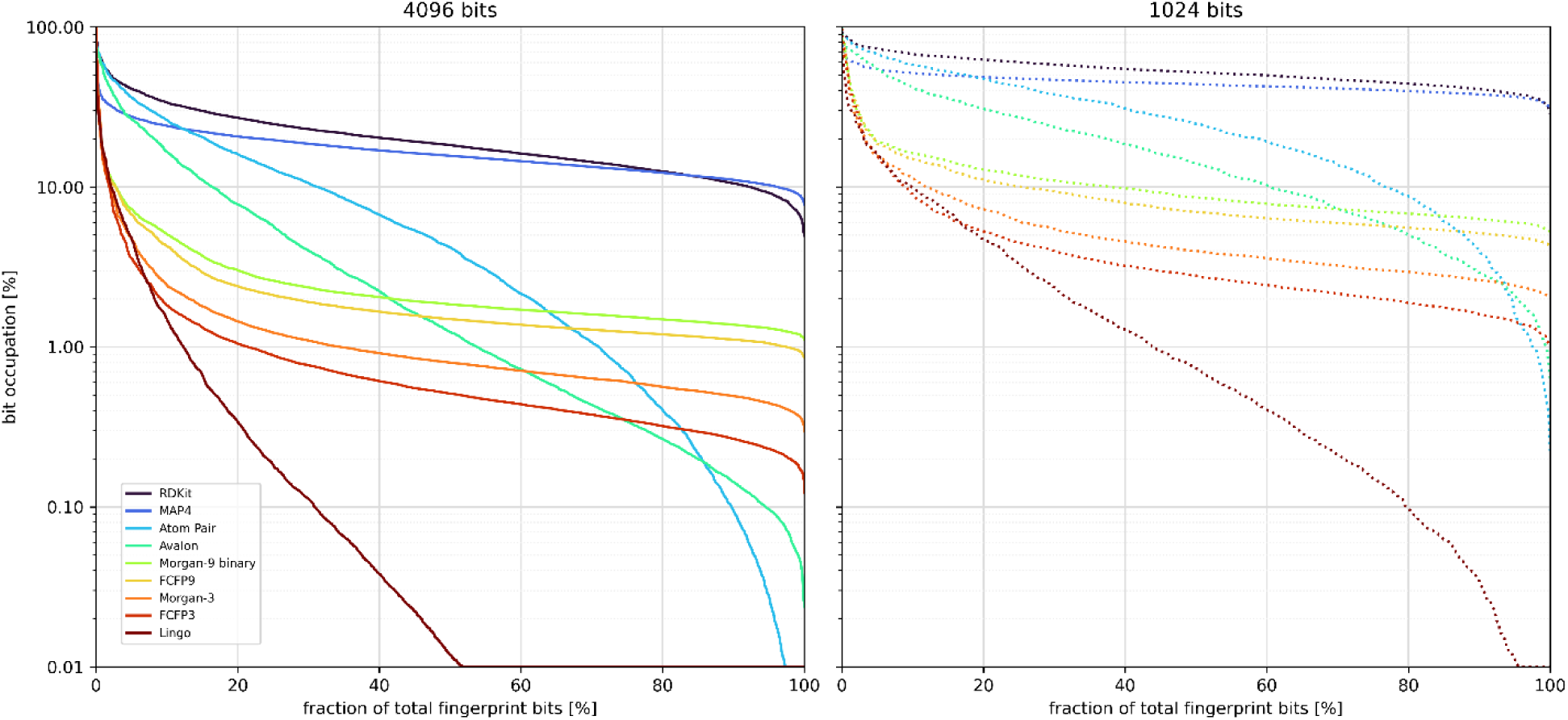
Bit occupation frequency for different molecular fingerprints displays a large variation for different fingerprint types. This ranges from nearly all bits being occupied > 30% of the time for 1024-bit RDKit and 1024-bit MAP4 fingerprints (dark blue and blue dotted lines in the right panel) to fingerprints with only a tiny fraction of bins occupied this often.

Switching to larger 4096-bit fingerprints strongly reduced the overall occupation rates for all fingerprint types, yet the large discrepancy between RDKit and Morgan or FCFP fingerprints remains. This reflects the more granular, local-neighborhood encoding strategy of Morgan-type, but also FCFP features.

MAP4, which encodes the distance between substructural motifs and thus captures a wide variety of fragment types, shows a bit occupation profile roughly comparable to the RDKit fingerprint, together forming the highest occupation fingerprints in our selection. On the other extreme end is LINGO with mostly very low bit occupations. Atom Pair and Avalon fingerprints show occupations rates between those extremes.

We here only measured the bit occupation which will not differ between count and binary variants of the selected fingerprints.

### Fingerprint duplicates

Because molecular similarity is inherently context-dependent, it is challenging to compare scoring methods in a strictly quantitative manner. Nonetheless, certain criteria can serve as clear-cut stress tests for any similarity measure. In line with the term “fingerprint”, one such property is the ability to avoid identical fingerprints for pairs of structurally distinct molecules.

We therefore identified all instances in which different molecules in the dataset shared an identical fingerprint. For each duplicated fingerprint, we computed the maximum mass difference between any two compounds bearing that same fingerprint. Figure 2 summarizes these statistics for both count and binary variants. The topological distance-based fingerprints, most notably MAP4, also Atom Pair, exhibited a particularly low number of duplicates, which indicates their ability to discriminate between highly similar compounds. For all other fingerprints, there is a large difference between binary and count variants with binary fingerprints showing more fingerprint duplicates. More importantly, however, is the strikingly large increase in duplicates with potentially very high mass differences often measuring tens of thousands of identical fingerprints between compounds with mass differences > 200 Da. Those compounds will naturally share key features, but they display large structural differences that intuitively should not result in identical vector representations (see examples in the supplemental material). Only Avalon fingerprints show similarly poor mass specificity in their count variant.

**Figure 2.**
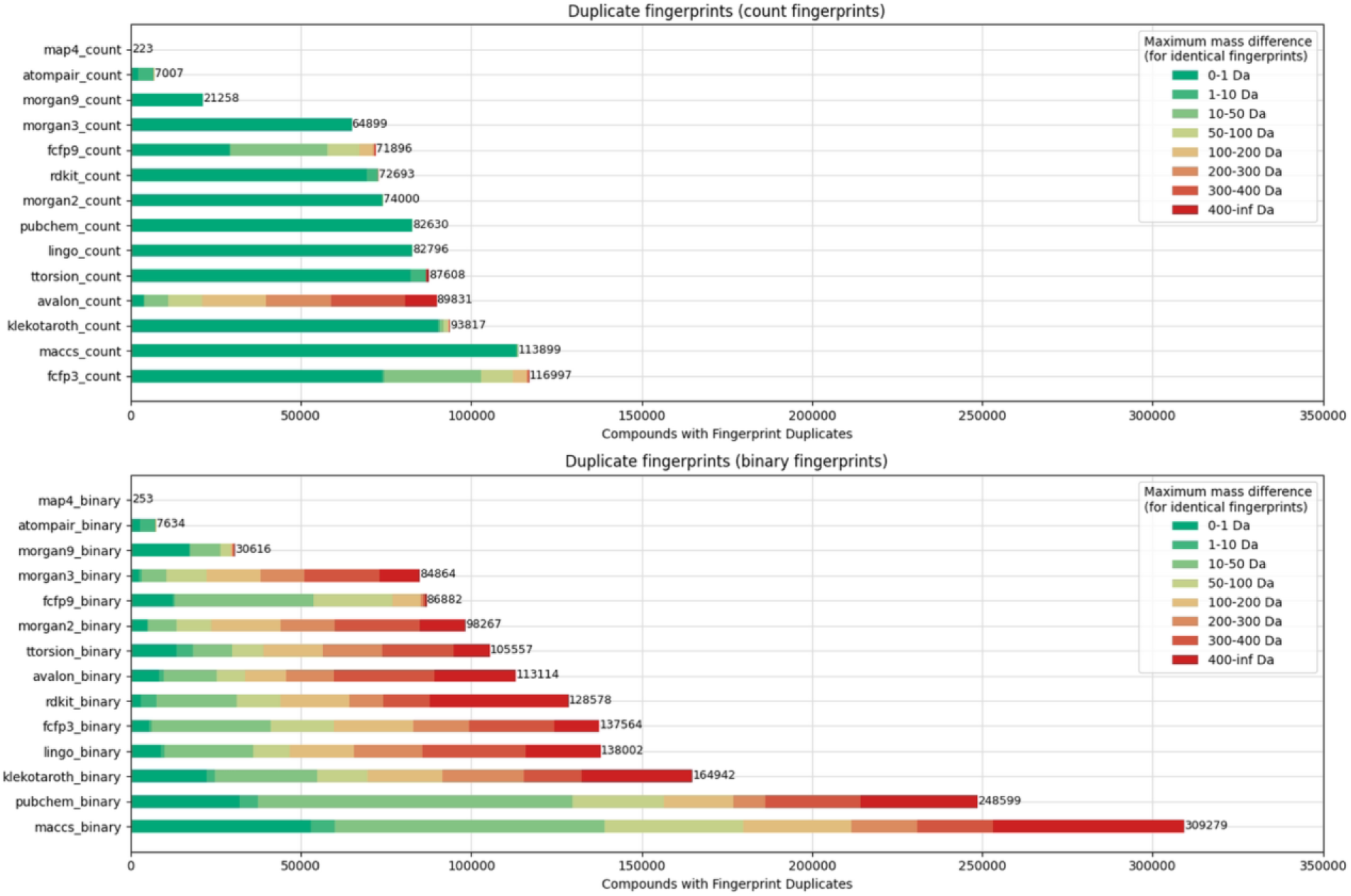
For different molecular fingerprints, all duplicates in a set of 718,067 unique compounds (biomolecular structures dataset) were counted. For each duplicate fingerprint, we computed the maximum mass difference among compounds sharing that fingerprint which is colored along different bins ranging from 0-1Da mass difference up to mass differences >= 400 Da.

Morgan and FCFP fingerprints exhibit an increase in specificity, indicating a reduction of duplicates with increasing radius. For Morgan fingerprints we observe a very notable drop from 64,899 to 21,258 duplicates (out of 718,067 unique compounds) when the radius is set from 3 to 9. Generally very low specificity can be seen for all dictionary-based fingerprints in our selection. In particular in their binary variant MACCS, followed by PubChem, and then Klekota-Roth fingerprints, all display very poor specificity. This improves when switching to count variants. Biosynfoni was only tested as count vector and displays duplicates for more than 50% of all compounds (see supplemental Fig. S1). A somewhat surprising observation was the performance of a simple element count vector as a baseline (see supplemental Fig. S1). As to be expected, this representation shows a very large number of duplicates (617,659), but nearly all of those are within a maximum mass difference of <= 1Da. This means that several fingerprints in our selection, FCFP3, FCFP9 and even more so Avalon and Biosynfoni, would already gain considerable specificity a simple concatenation with a 11-dimensional element count vector.

### Fingerprint Comparisons Reveal Divergent Perspectives on Molecular Similarity

Different fingerprint representations capture distinct aspects of molecular structure, and these differences are reflected in their similarity score distributions. To give a first intuition, we here compare RDKit fingerprints and Morgan fingerprints, across the 37,811 compounds in the **ms2structures dataset**. Although both fingerprint types are commonly employed in tasks such as compound ranking, their underlying algorithms and bit occupancy rates differ noticeably, leading to different interpretations of what it means for two molecules to be similar.

Due to its higher bit occupation rates, the default RDKit fingerprints generally produce higher Tanimoto similarity scores compared to Morgan-3 fingerprints. To enable a more meaningful comparison, the raw similarity scores were converted into percentile values ranging from 0% to 100% based on all unique compound pairs (see methods). This scaling allowed us to directly contrast how each fingerprint representation ranks molecular similarity across the entire dataset.

#### Comparison 1: Morgan-3 Binary vs. RDKit Binary Fingerprints

When comparing Tanimoto similarity scores computed from Morgan-3 binary (4096-bit) fingerprints with those derived from RDKit binary fingerprints of the same bit length, substantial differences in their percentile distributions were observed. Notably, a significant proportion of compound pairs, approximately 43.41%, were consistently assigned to the bottom 60% of similarity by both scoring methods.

In contrast, only a smaller fraction of pairs (0.42%) was jointly classified within the top 1% of similarity. Moreover, there were striking discrepancies for certain compound pairs. For instance, over 200,000 pairs (0.04% of all unique pairs) were placed in the top 1% by RDKit fingerprints while being relegated to the bottom 60% by Morgan-3 fingerprints.

Conversely, more than 900,000 pairs (0.13%) were ranked in the top 1% by Morgan-3 fingerprints but fell into the bottom 60% based on RDKit scores (Figure 3). These results underline that even when using the same Tanimoto metric, the two fingerprint types emphasize different molecular features, resulting in divergent similarity assessments.

**Figure 3.**
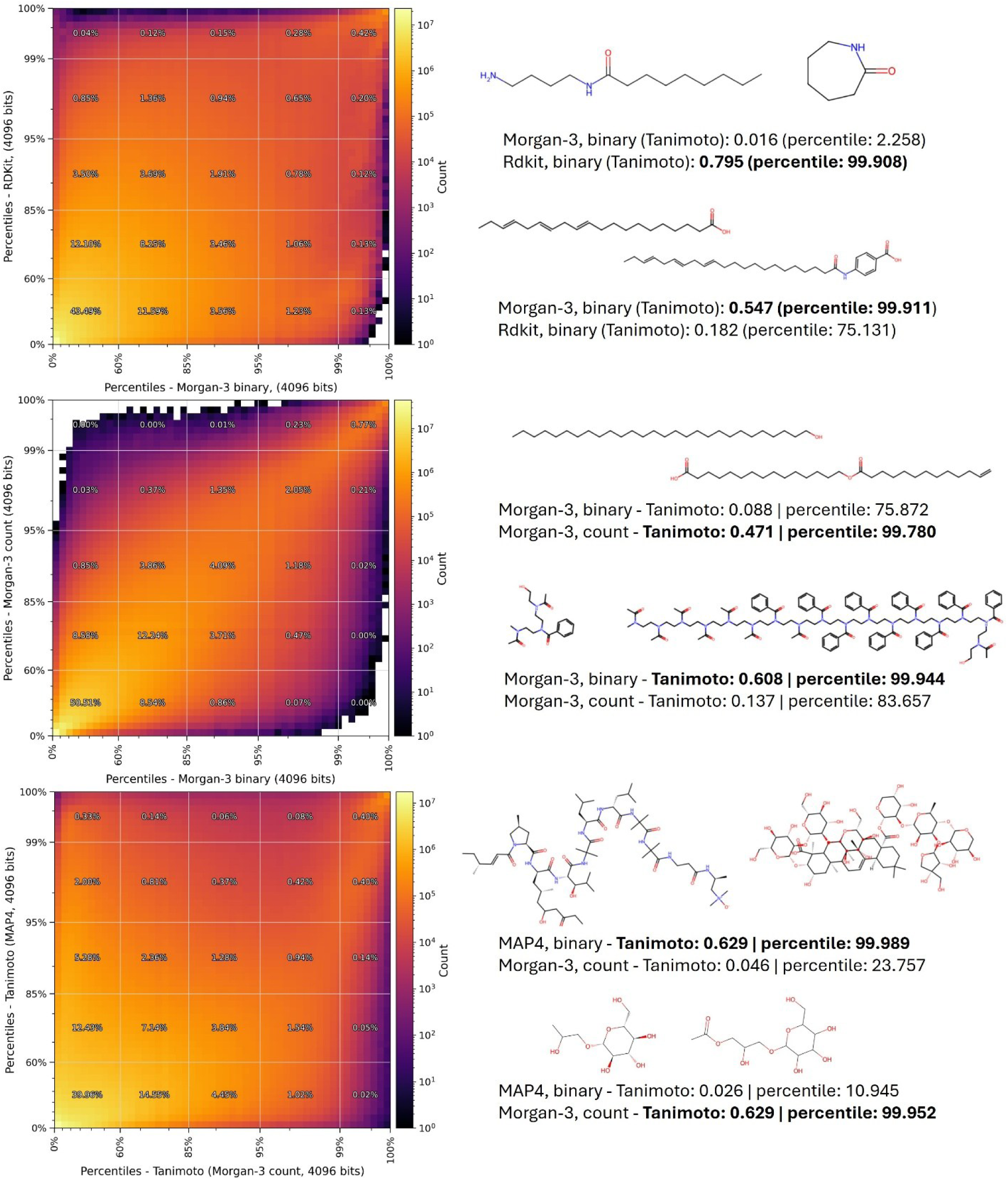
Comparison of all pairwise similarities for the ms2structures dataset (37,811 compounds) using various fingerprints. **(top row)** Tanimoto scores computed from RDKit fingerprints and Morgan-3 binary fingerprints. **(center row)** Tanimoto scores computed from Morgan-3 binary fingerprints and from Morgan-3 count-based fingerprints. **(bottom row)** Tanimoto scores were computed from Morgan-3 count fingerprints and from MAP4 fingerprints. All fingerprints were 4096-bit vectors. Example pairs with drastic score discrepancies are highlighted on the right.

#### Comparison 2: Morgan-3 Binary vs. Morgan-3 Count

To further probe the influence of fingerprint representation, we compared Tanimoto scores computed from Morgan-3 binary fingerprints with Tanimoto similarity scores derived from Morgan-3 count-based fingerprints. In this pairing, both methods are based on the same underlying fingerprint type, yet the representation (binary versus count) introduces noteworthy differences.

The analysis revealed that about 50.42% of compound pairs were consistently ranked in the bottom 60% by both metrics, while 0.77% of pairs were jointly assigned to the top 1%. Although the overall agreement between the two metrics is higher than in the previous comparison, notable differences persist for certain compound pairs.

The count-based fingerprints offer the advantage of recognizing repeated occurrences of motifs or substructures, as exemplified in an example pair in Figure 3, providing a more nuanced view of molecular similarity. However, this sensitivity can also be a drawback, as it may overemphasize the contribution of common substructures, such as extended carbon chains (e.g., CC chains) leading to disproportionately high similarity scores in some cases.

#### Comparison 3: MAP4 vs. Morgan-3 Count

The third comparison involves Tanimoto similarity scores computed using MAP4 fingerprints and from Morgan-3 count-based fingerprints, with both representations using 4096-bit vectors. As illustrated in Figure 3 (bottom row), the overall correspondence between these two metrics is less pronounced than in the previous comparisons. Only 39.97% of compound pairs fall into the bottom 60% for both metrics, and just 0.39% are classified in the top 1% by both methods.

Notably, MAP4 exhibits the highest bit occupation among the fingerprints evaluated, a characteristic that appears to lead to a substantial number of bit collisions, particularly for larger molecules. This increased collision rate manifests in a sizable subset of cases: approximately 0.33% of pairs are ranked in the top 1% for MAP4 while simultaneously being assigned to the bottom 60% according to the Tanimoto score on Morgan-3 count fingerprints.

In addition, a smaller group of pairs is observed in the opposite corner of the score distribution, where the Morgan-based count fingerprints place them in the top 1% while MAP4 assigns them to the bottom 60%. We noted that those pairs are usually clear cases of unambiguous false similarity assignments by MAP4 as indicated by the two example pairs in Figure 3.

### Unfolded fingerprints to avoid bit collisions

Molecular fingerprints are usually brought to a fixed size. Common in literature, tutorials, and library default settings are vector sizes of 1024, 2048, or less frequently up to 4096 bits.

Folding the found substructure bits into fixed-sized vectors is an efficient trick to simplify further use of such fingerprints, both conceptually as well as computationally. However, this approach results in information loss, and bit collisions are a well-known downside. When the number of bits is high enough, the effect of bit collisions on ranking tasks was described to be very low by some ^1^ while others reported that, in particular for very close analogues, results improved notably when using 16384-bit fingerprints instead of 1024-bit ones ^62^. However, due to the focus on very close analogues, this might not fully reveal the extent to which bit collisions might influence the all-vs-all comparisons of a larger number of molecules.

A large part of the very pronounced differences between the various similarity scores as shown in Figure 3 can be attributed to fundamentally different fingerprint concepts. Yet, at the very least a notable fraction of very high discrepancies between MAP4 and Morgan-3 based fingerprints pointed at more substantial fingerprint-related shortcomings (see pair examples in Figure 3). Consistent with the high overall bit occupation ratio shown in Figure 1, this suggests severe effects from bit collisions. Consequently, we also computed unfolded fingerprints of all tested variants, which kept all original bits.

We computed Tanimoto scores for all unique pairs between the 37,811 compounds in the ms2structures dataset using both unfolded as well as 4096-bit Morgan-3 count fingerprints. The results confirm that the scores are generally only very mildly affected (Figure 3, left).

Interestingly, in the case of Morgan-3 count fingerprints, we found that computing molecular similarities for unfolded fingerprints was about five times faster than for Morgan-3 folded 4096-bit fingerprints. This can be explained by a relatively low number of bits per molecule for Morgan-3 (see Figure 5). For fingerprints with a much higher number of occupied bits, such as the RDKit fingerprint with the default setting of paths up to the length 7, handling unfolded fingerprints is still feasible but with notably slower similarity score computations.

For the RDKit fingerprints, as well as for folded MAP4 fingerprints, unfolded fingerprints lead to very pronounced shift in similarity scores, shown in Figure 4. Both for RDKit and MAP4 fingerprints, this reveals a very large fraction of pairs that have little to no overlap in actual bits (as represented in the unfolded vectors), but which are shifted towards values in the 0.1 to 0.2 score range due to the folding of bits to a fixed vector size of 4096. Effectively, this raises the average similarity scores for such fingerprint types.

**Figure 4.**
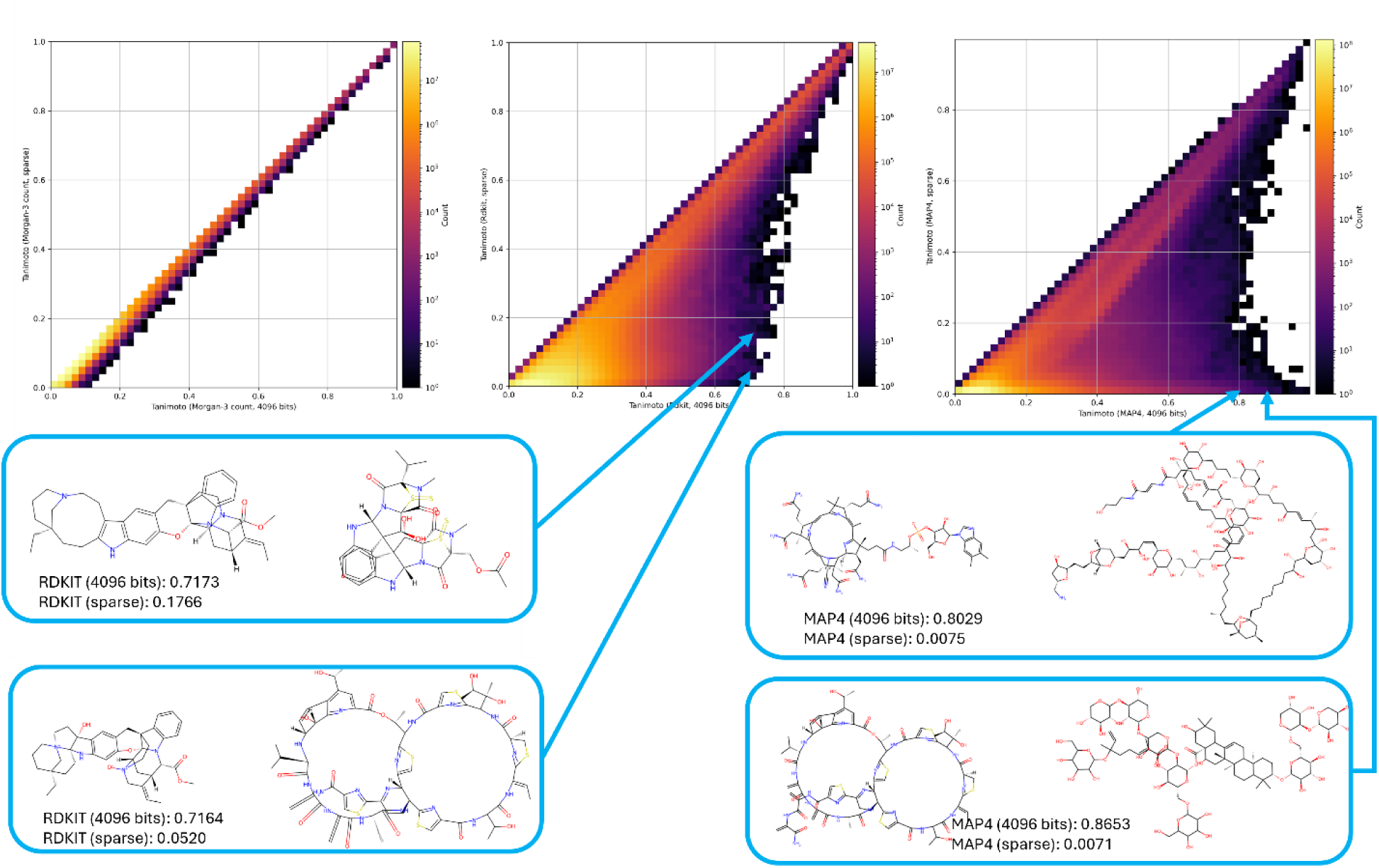
Comparison of unfolded vs folded 4096-bit RDKit fingerprints on all possible unique pairs between 37,811 compounds (ignoring pairs between identical compounds). Shown are Tanimoto scores for Morgan-3 count fingerprints (left), Tanimoto scores for RDKit fingerprints (center), and Tanimoto scores for MAP4 fingerprints (right). Due to bit collisions in the RDKit and MAP4 fingerprints, the scores based on 4096-bit vectors are generally notably higher. In some cases, pairs that should have a very low similarity according to the respective fingerprint algorithm do receive very high scores due to bit collisions (see example pairs in the blue boxes).

**Figure 5.**
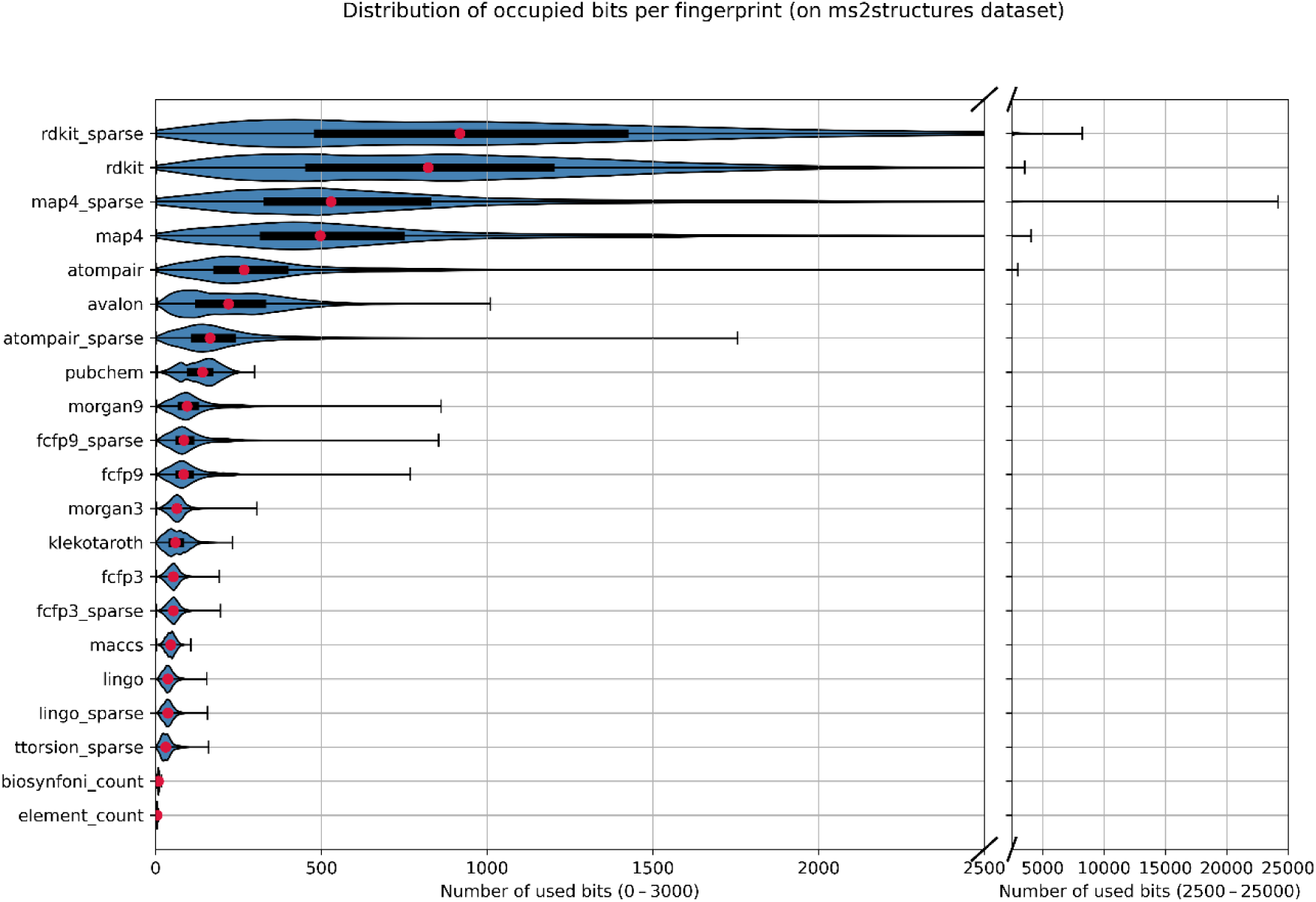
Distributions showing the number of non-zero bits over the entire ms2structures dataset. The subplot on the right shows the long tail of some of the distributions towards reflecting few compounds which led to very high numbers of occupied bits. The unfolded MAP4 implementation, for instance, resulted in fingerprints of up to 24,183 bits.

In addition, a smaller but notable number of pairs erroneously receive very high similarity scores of up to 0.6 or higher, even though they correctly belong in the 0 to 0.2 score range. The latter is particularly dangerous for ranking or search tasks because this can lead to arbitrary false hits.

### Compound size dependence

The number of possible substructures of a compound generally increases with increasing compound size. Therefore, heavier organic compounds should, on average, have a higher number of occupied bits in a molecular fingerprint, as well as higher values for count vectors. It has been shown that this difference in bit occupation can result in different statistical properties of similarity scores ^63^.

Here we compare all similarity scores between the smaller molecules in the ms2structures dataset (< 300 Da, 8,601 compounds, 36,984,300 pairs) with all scores between larger molecules (> 500 Da, 8,570 compounds 36,718,165 pairs). Both groups contain an approximately equal number of compounds to avoid effects based on the sample size. Out of the scores for all possible pairs in each group, score percentiles starting at 50% were computed and are displayed in Figure 6.

**Figure 6.**
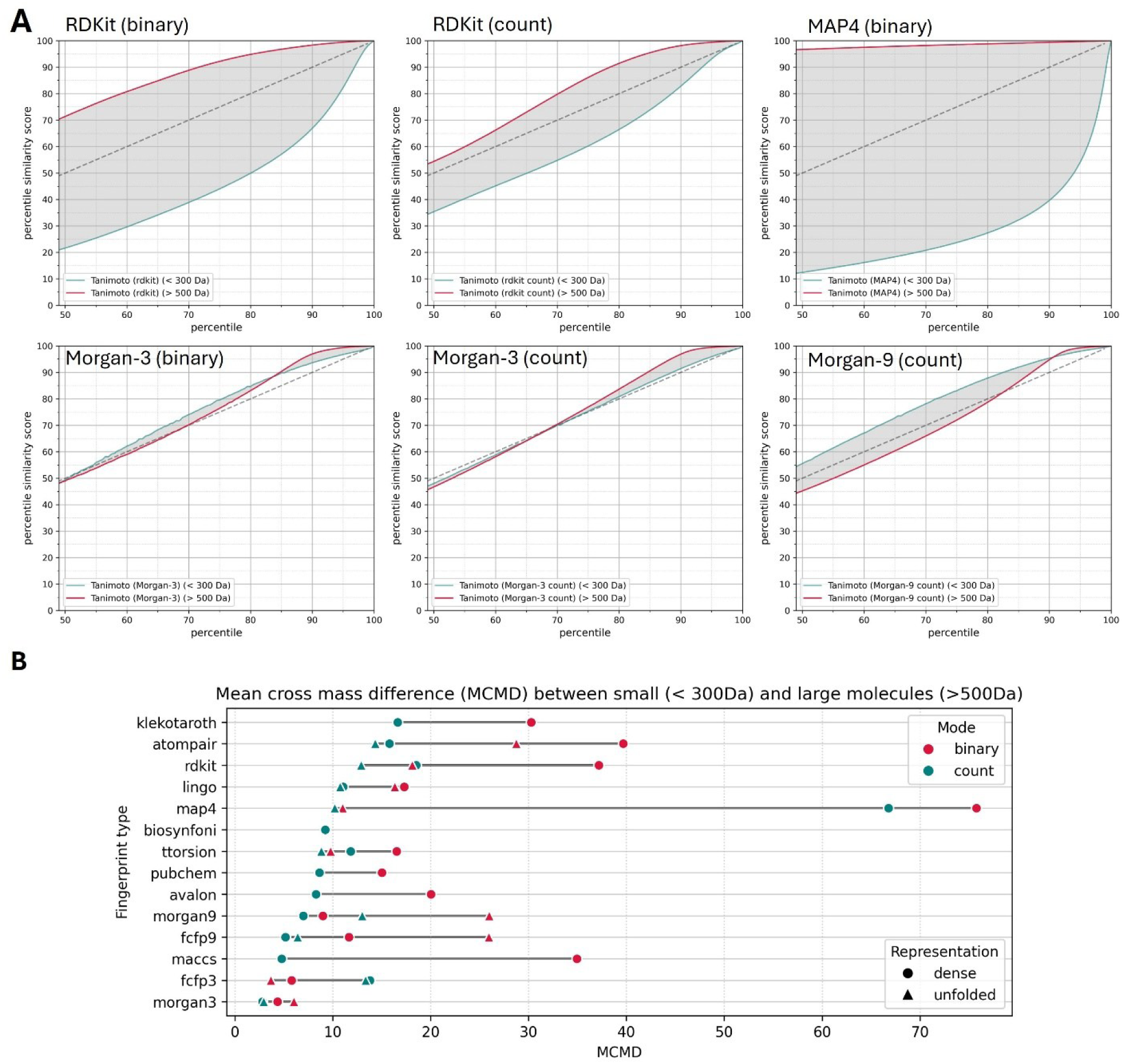
**(A)** Fingerprint-based similarities are computed for about 36 million unique pairs between small compounds (< 300 Da) or between larger compounds (> 500 Da). The similarity scores of increasing percentiles are plotted for both the small and larger molecules, ranging from 50% up to 99.5%. This is done for Tanimoto scores of RDKit fingerprints (top left), RDKit count fingerprints (top center), MAP4 fingerprints (top right), Morgan-3 binary fingerprints (bottom left), Morgan-3 count fingerprints (bottom center), as well as Morgan-9 count vectors (bottom right). All fingerprints used had 4096 bits. **(B)** The mean difference between the high and the low mass curves was computed for multiple fingerprints, both using a folded and unfolded implementation (4096 bits, except for dictionary-based fingerprints: MACCS, PubChem, Klekota-Roth, and Biosynfoni).

In Figure 6A we exemplarily display the percentile similarity score curves for some RDKit, MAP4 and Morgan-3 and 9 variants. For the fingerprints with high bit occupation frequencies, RDKit and MAP4, the similarity score percentiles are consistently much higher for larger molecules.

For the tested Morgan-3 fingerprints, this is only the case for higher percentiles starting at about 85 - 90% and the effect generally remains moderate. The median of all similarity scores, which is the percentile at 50%, is hence largely the same within small and larger molecules in the case of Morgan-3 fingerprints, both for binary fingerprints and count fingerprints.

In contrast, for RDKit and MAP4 fingerprints, the median score varies considerably basedon the molecular mass. For instance, for the RDKit binary fingerprint, about 15% of all pairs between larger molecules (> 500 Da) received a similarity score which was higher than 0.3, thus placing it in the top 2% of all scores. At the same time, only about 1% of pairs between smaller molecules (< 300 Da) received similarly high scores (see also Figure S1 in the supplemental material).

To better compare many different fingerprints, we computed the mean difference between the high and the low mass curves, which we will refer to as the mean cross-mass difference (Figure 6 B). This allows several general observations. With the exception of FCFP3, all fingerprints show lower cross-mass differences in their count variant. This is particularly notable for MACCS, Atom Pair, Klekota-Roth, RDKit, and Avalon fingerprints.

The effect of moving from folded to unfolded fingerprints is less consistent. MAP4 improves drastically when left unfolded, and also Atom Pair and RDKit, followed by topological torsion fingerprints show a notable decrease in cross-mass difference. Interestingly, though, we see higher mean cross-mass differences for similarity scores computed for unfolded Morgan-9 or FCFP-9 fingerprints when compared to fixed-size 4096-bit vectors.

### Focus on the top hits: Ranking

The score comparisons over the entire dataset illustrate that the different scores and fingerprints can lead to sometimes drastically different interpretations of similarity. For certain downstream tasks the full range of scores is of high interest. For instance, for training machine-learning models to predict chemical similarities (e.g., MS2DeepScore ^50,64^) or models trained to predict compound structures which are then evaluated based on chemical similarity measures, typically Tanimoto scores of common molecular fingerprints ^7–9,65^.

For other tasks, such as nearest neighbor searches, however, one might argue that it is of little importance if a pair of compounds is in the lowest 20% percentile or in the lowest 50% percentile since the entire focus is set on finding the most similar compounds. The direct score comparisons in Figure 3 display that even in the highest percentiles the scores occasionally differ, but this is probably better assessed using top-n rankings. Unlike virtual screening tests such as Riniker et al ^1^, we here first want to inspect the ranking agreement between the different scores across a random subsample of 10,000 compounds from the ms2structures dataset.

For each of the 10,000 compounds, the closest 10 neighbors are computed for all fingerprint types and variants used in this work. To account for potential issues due to bit collisions, both 4096-bit and unfolded fingerprints were used where feasible. In the next step, the overlap in the top-10 selections was measured between all possible score combinations to compute the mean top-10-overlap. The full cluster-ordered heatmap showing all top-10-overlaps between all of the 45 used fingerprint types and variants can be found in the supplemental material.

Consistent with the very moderate difference between folded and unfolded variants of Morgan, FCFP, or LINGO type fingerprints, we see a relatively high selection agreement of, on average, more than 9 out 10 candidates. This value is lower for RDKit binary fingerprints (8.3) and most pronounced for the binary variants of topological torsion (6.58), MAP4 (6.64), and Atom Pair fingerprints (6.67).

To better illustrate the 45 different types and variants, we computed a minimum spanning tree connecting types and variants by their respective top-10-overlap values (Figure 7). This also displays a clear separation between the respective binary and count variants. For MACCS, for instance, this shows only a top-10 overlap of 3.76 essentially making its top-10 count-based ranking more consistent with Atom Pair count fingerprints than with its own binary counterpart. In general, this ranking comparison reveals an often low agreement in what different fingerprints will select as most-similar compounds with a majority of cases having top-10 overlaps below 4.0 out of 10 (median is 3.46, mean is 3.76, see heatmap in supplemental material).

**Figure 7.**
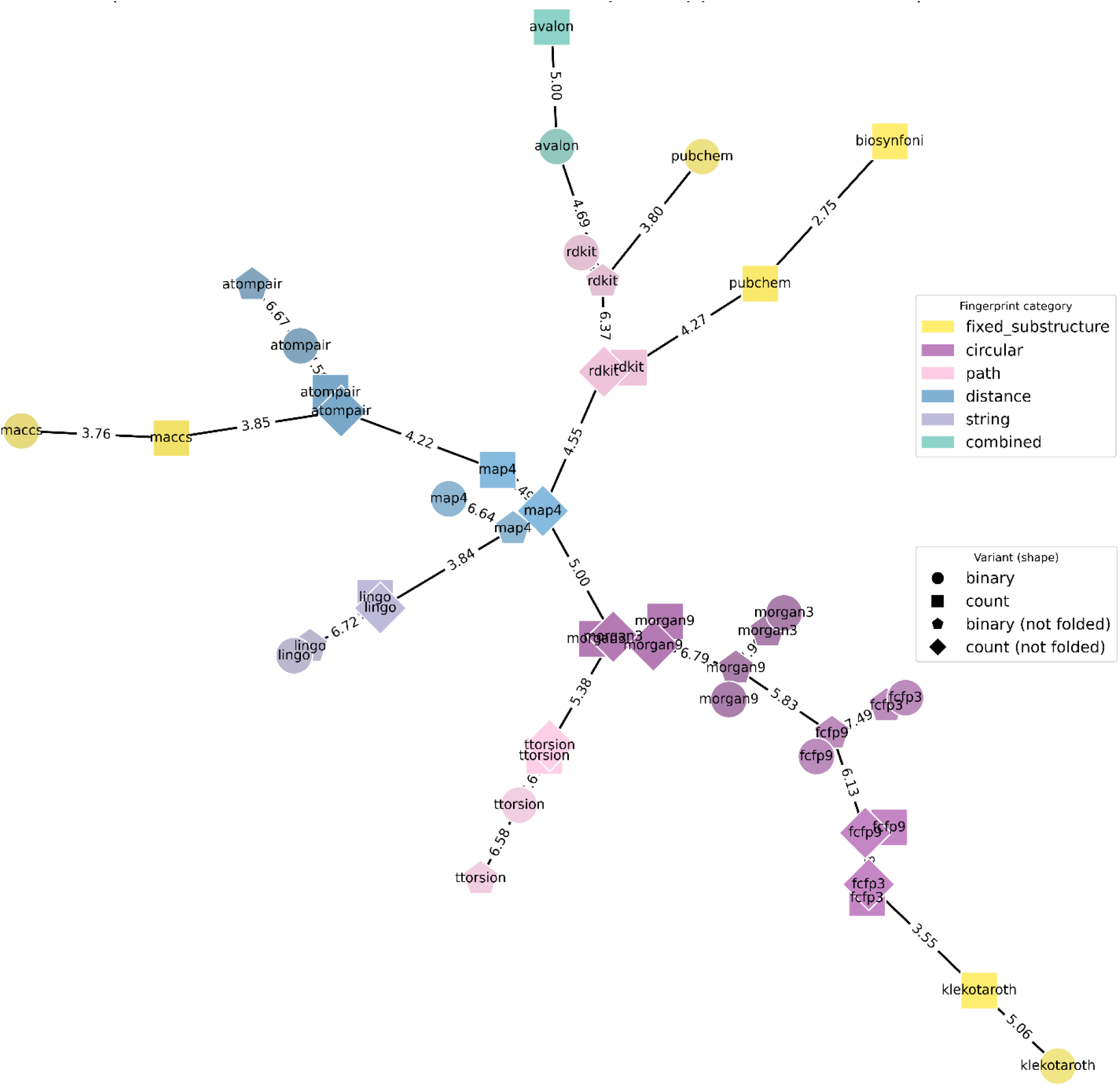
For a random subset of 10,000 compounds from the ms2structures dataset, the ten most similar compounds were selected based on the different fingerprints. For all possible combinations, the average overlap between all top-10 selections was computed and used to generate a minimum spanning tree of all fingerprints based on their respective mean top-10 overlaps. Each node in this graph represents one fingerprint variant. The shape of each node represents the variant (binary, count, and folded/not folded), the node colors represent the fingerprint category. Numerical values on the edges stand for the respective mean top-10 ranking overlaps between adjacent nodes.

Ranking compounds is key in many large-scale searches. Such tasks in the form of virtual screenings are probably the most common approach to benchmark molecular fingerprints and similarity measures with the open-source benchmarking platform of Riniker and Landrum ^1^. This has recently been expanded by tasks involving peptides ^14^. MAP4 fingerprints, and more recently chirality aware MAP4c fingerprints ^15^, were shown to perform particularly well on this benchmarking set. In the absence of peptides, the superior average ranking of MAP4 is often not statistically significant, including when compared to Morgan-2 2024bit binary fingerprints (see supplemental material in ^14^).

Here we do not want to fully repeat the Riniker and Landrum virtual screening task (see more discussion and exemplary tests in the supplemental material). Instead, we aim to assess the fingerprint’s ability to predict various activity classes based on the work of Boldini et al. ^17^. They used data on 12 different bioactivity classes to train 12 different classifiers. We slightly adjust this task by combining all bioactivity labels in one dataset, including compounds from their dataset which do not belong into any of those classes (see methods). Since compounds can belong into more than one class, this creates a multi-label classification task which is slightly more complex than the original setting ^17^. While this might not guarantee the best possible performance for each individual class, this altered setting will only require the training of one model for all classes at the same time, making this a more efficient benchmarking setup. One might also argue that this setting is maybe closer to reality since conflicting labels are a common phenomenon.

For each fingerprint variant, a shallow neural network model was trained as a classifier for all 12 bioactivity classes (see methods). Since the (initial) training of such models can be hampered by high inputs, we always scaled the counts logarithmically, which means that we cannot differentiate between count and log-count variants in this experiment.

While all fingerprints allowed to reach considerable accuracies for the 12 classes (Figure 8), the high label bias in the dataset meant F1 scores were consistently lower. Judging by the macro F1 score, MAP4 and FCFP9 resulted in the best performing classifiers. Given the small number of labels for some classes, sometimes as low as 20 or 22 samples per class, the variability in performance results in difficulty in clearly ranking the subsequent fingerprint types such as Atom Pair, Morgan, or RDKit. Fairly characteristic, however, is that all dictionary-based fingerprints (MACCS, Klekota-Roth, PubChem, Biosynfoni) generally resulted in lower model performance.

**Figure 8.**
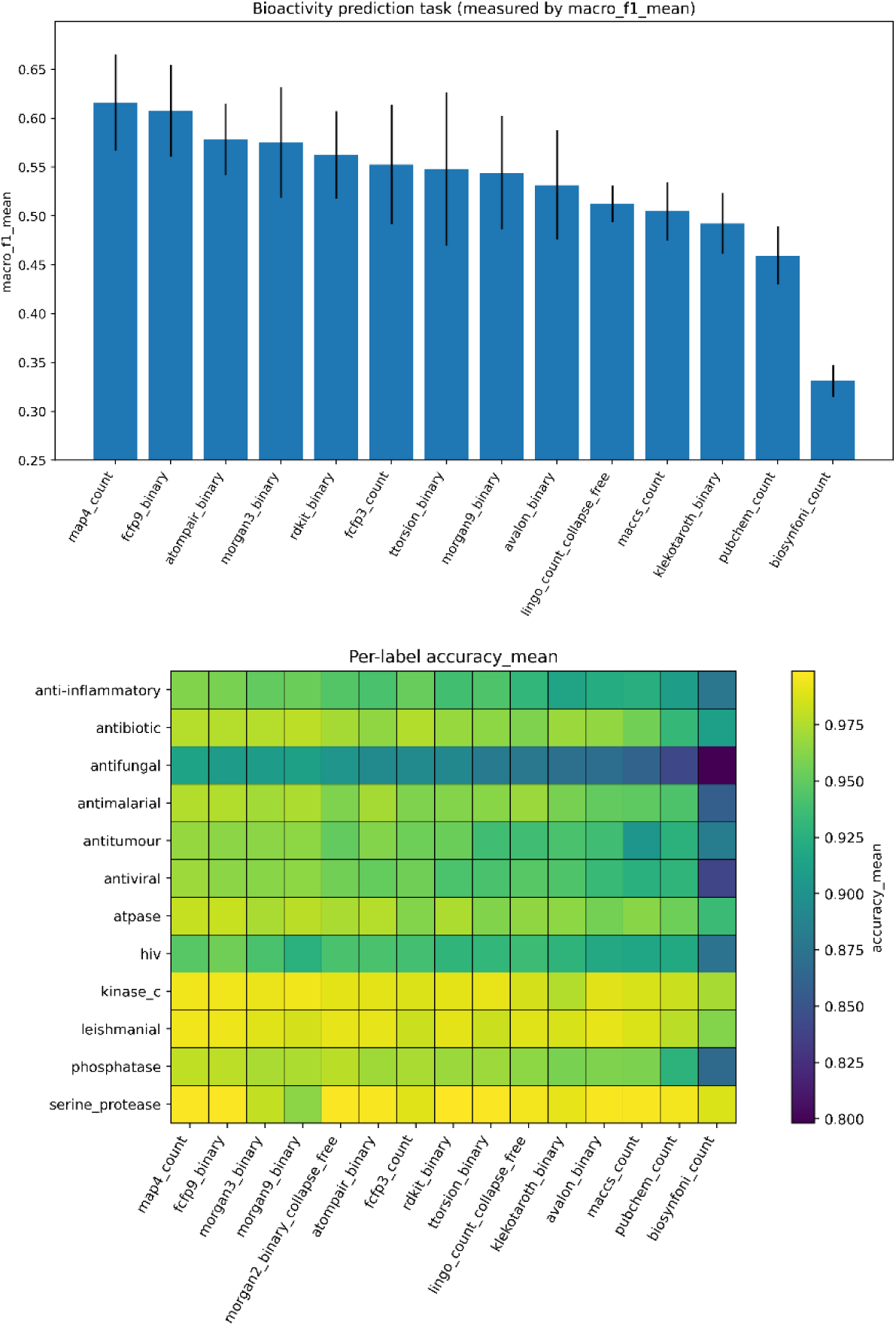
Simple deep-learning models (shallow neural networks) were trained on predicting 12 different bioactivity classes based on various fingerprint types and variants. This was done using a 5-fold split to also assess the performance variability. For each fingerprint type, the best performing variant was selected (from binary, count, as well as frequency-folded binary and frequency-folded count). Their resulting macro F1 is shown in the upper bar plot with error bars indicating the variation among the 5-fold splits. (Bottom) The average accuracies are displayed for all fingerprint types (again, selecting the best variant) and all 12 bioactivity classes.

Another observation was that for each specific fingerprint type, the variant choice (binary or count) hardly affected the model performances, thereby indicating that the presence or absence of specific substructures seems to be enough for a machine-learning model to predict the bioactivity class.

### Comparison to graph-based method

As described in the introduction, there is no generally valid ground truth chemical similarity measure. A conceptually compelling approach, however, is graph-based measures to assess the maximum common subgraph (MCS) or maximum common edge subgraph (MCES) of two compounds.

Those measures come in different flavors and implementations, but all share the disadvantage of a very high computational cost when compared to the above use of fingerprint-based methods. Fingerprint computations scale very favorably with molecular size, and once fingerprints are computed, the actual computation of molecular similarities is compound-size-independent, at least for fixed-sized vectors.

In contrast, MCS/MCES measures can sometimes require substantial computational resources for only a few such calculations. Even when making use of lower bound estimates, this disadvantage can only partly be mitigated ^6,37,38^. In addition, such optimization by design makes the scores less suitable for computing chemical similarities between dissimilar compounds.

We here computed RascalMCES ^37^ scores implemented in RDKit ^32^ for a sampled subset of 5,413,677 compound pairs out of the ms2structures dataset (see Methods). To avoid too many trivial cases of entirely unrelated molecules, we only sampled from pairs with a mass difference of up to 100 Da. The rascalMCES scores were then compared to the different fingerprint-based similarity measures by computing the Spearman correlation. This measures how often a higher fingerprint-based similarity corresponds to a higher MCES similarity. As a baseline we also tested a simple 11-dimensional element count vector in this experiment.

For all fingerprints for which we tested both folded and unfolded variants, the unfolded variants show better correlation values (Figure 9). For some fingerprints such as FCFP3 or LINGO this is only a minor effect, but Atom Pair, RDKit, and in particular MAP4, all improve very notably when switching to unfolded variants. In addition, throughout the fingerprint types, it can be observed that count variants correlate better with rascalMCES scores. The exceptions are FCFP3 and LINGO fingerprints for which this effect is small to nonexistent.

**Figure 9.**
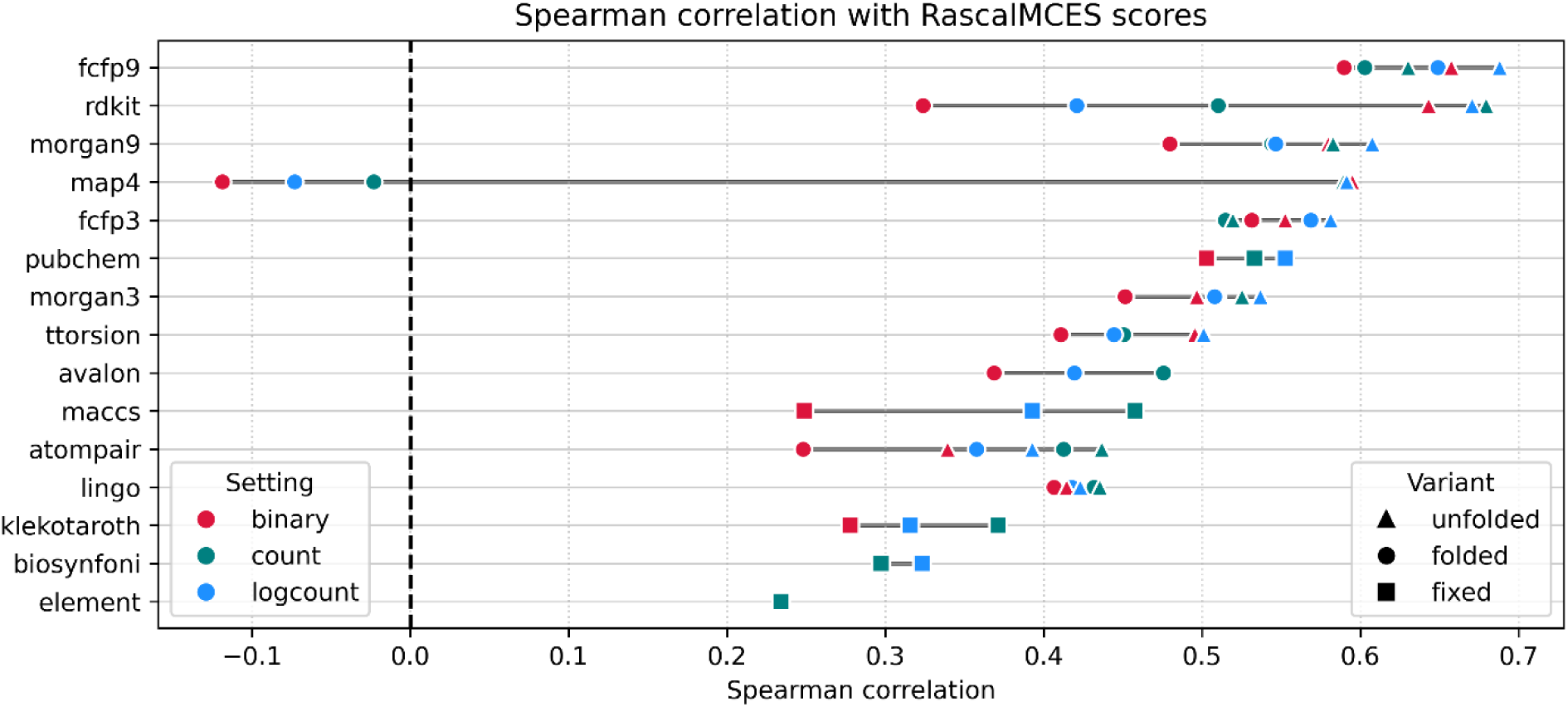
Spearman correlation between rascalMCES scores and various fingerprint-based similarity measures. For each fingerprint, both 4096-bit (red circles) and sparse (teal squares) implementations were used. The correlation coefficients were computed based on 5,413,677 randomly sampled compound pairs from the ms2structures dataset (see methods).

Considering that a simple element count baseline already reaches a Spearman correlation of about 0.23, several fingerprint types such as Klekota-Roth, Biosynfoni, and in their binary folded variants also MACCS, Atom Pair, and RDKit fingerprint, show rather poor correlation values.

MAP4 is a special case here, because for all its folded variants show practically no overall correlation with rascalMCES scores.

The latter is in sharp contrast to the unfolded MAP4 fingerprints variants, which reach relatively high Spearman correlations of about 0.59. Only Morgan-9 (unfolded log count), RDKit (all unfolded variants), or FCFP9 (virtually all variants) reach better correlations values of up to about 0.69 (FCFP9, unfolded log count).

When comparing simple integer counts with log counts, our results do not allow a general recommendation. While the log count variants slightly outperform other variants for Morgan, FCFP, or PubChem fingerprints, we see an opposite effect in other fingerprints such as MACCS, Avalon, or Atom Pair.

### Chemical class prediction

Molecular fingerprints cannot only be used as input for models to predict bioactivity, but also to predict taxonomic origin ^47^ or chemical classes ^16,25,26^. Unlike work on creating reliable classifiers ^25,26^, the main goal here was not to create the most performant classifier model. Instead, our focus was to evaluate each fingerprint’s potential to encode the required information for such classifier models. We therefore selected a relatively simple shallow neural network architecture and used a balanced subset containing 1000 compounds for each of 120 different chemical subclasses as defined using Classyfire ^66^.

We again compared binary and count variants of each fingerprint type. Unfolded fingerprints are not feasible for this task since the trained model requires a fixed sized input with a manageable number of input dimensions. To still test if folding has a relevant impact on the model performance, we also used a frequency folded variant by only selecting the 4096 most occupied bits from an unfolded variant, thereby avoiding any bit collisions.

A relatively simple network model was used to predict all 120 subclasses from a fingerprint input (see methods). As with the bioactivity prediction task, we scaled the counts logarithmically and can therefore not differentiate between count and log-count variants in this experiment.

For all fingerprint types, simple neural network models trained on count variants perform notably better than models trained on binary variants, with the largest shift for MACCS fingerprints, but also the other dictionary-based fingerprints (PubChem and Klekota-Roth), see Figure 10.

**Figure 10.**
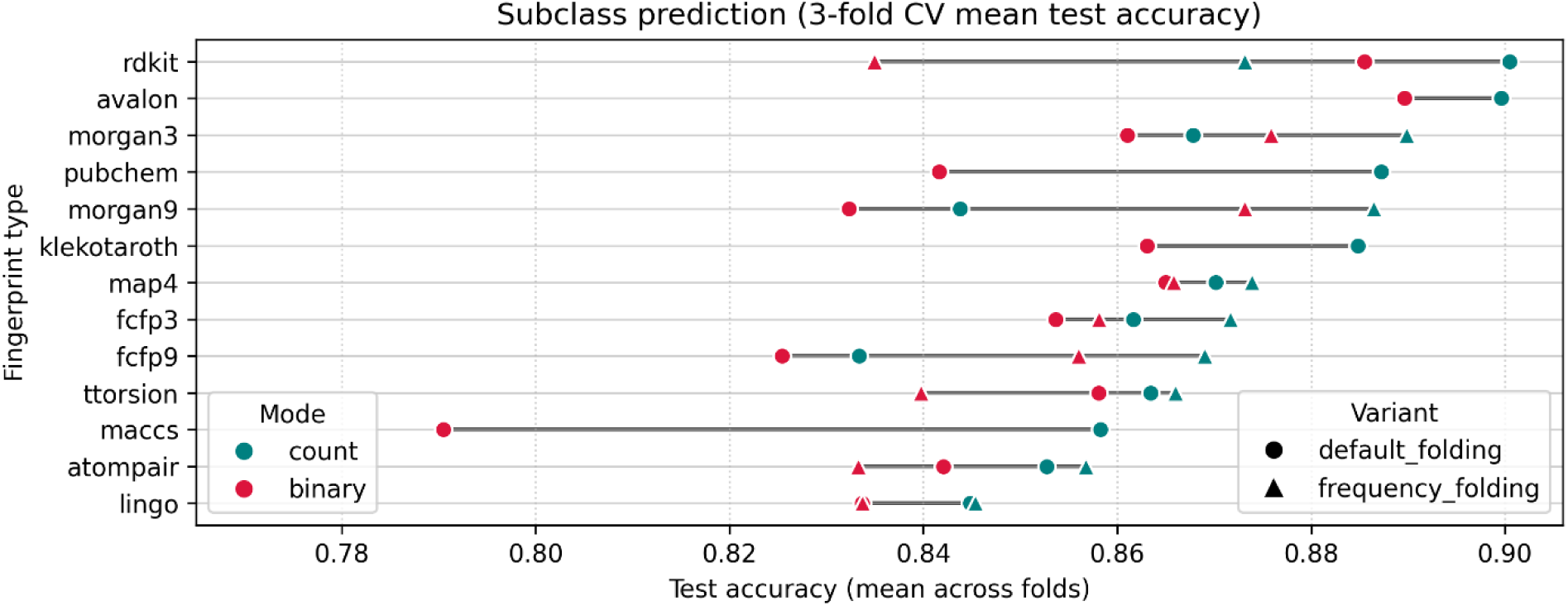
Average accuracy on the balanced 120-subclass dataset for subclass prediction. Except for the fixed-sized substructure fingerprints (PubChem, MACCS, Klekota-Roth) all fingerprints were brought to 4096 bits either by standard folding (default folding) or by selecting the 4096 most occupied bits over the entire dataset (frequency-folding) for fingerprints that provided an unfolded variant.

The effect of the folding was more dependent on the fingerprint type. A larger performance gain was observed for Morgan and FCFP fingerprints, in particular Morgan-9 and FCFP9. For MAP4 we only observe a negligible effect, and for Atom Pair and torsion fingerprints we see a small to negligible gain for the count variant when switching to frequency folding, but a larger performance loss for the binary variant. Most surprising to us was the very notably drop in performance for RDKit fingerprints upon switching to frequency-folded variants.

Overall, RDKit fingerprints and Avalon fingerprints performed best on this task, followed by Morgan, PubChem, and Klekota-Roth fingerprints. Biosynfoni is not shown in Figure 10 since its performance was not on par on this classification task (see supplemental material).

### Chemical class consistency

Training subclass classifiers aimed at assessing how much required information fingerprints contain for predicting chemical classes. A related, yet different perspective is to look for chemical class (or subclass) consistency within the abstract vector space described by each fingerprint type and variant. It can be expected, that compounds of the same chemical subclass, here again defined using Classyfire ^66^, will frequently be close to each other. Defining chemical classes or subclasses, is an extremely complex task, not least due to a large variety of origins, rules, and conflicting overlapping categories ^6,26,66^. Therefore, such labels cannot be regarded as sharply defined ground truth. It is, for instance, chemically plausible that a compound may be found near a different compound belonging to another class or subclass which at times might only be rooted in a minor modification in their overall molecular structure.

Nevertheless, chemical classes are well-suited to represent overarching chemical properties, which is why they are also often used in more complex visualizations of chemical spaces where they indeed often display a high spatial homogeneity ^6^.

To assess subclass consistency, 3000 compounds of each of the 25 most frequent subclasses in the biostructures dataset were selected, giving a subclass balanced dataset of 75,000 compounds in total (see methods). The 100 nearest neighbors of each compound were computed using an approximate nearest neighbor search using the inverse Tanimoto similarity as a distance measure (see methods). We then computed the fraction of same-subclass compounds within those 100 nearest neighbors, see Figure 11.

**Figure 11.**
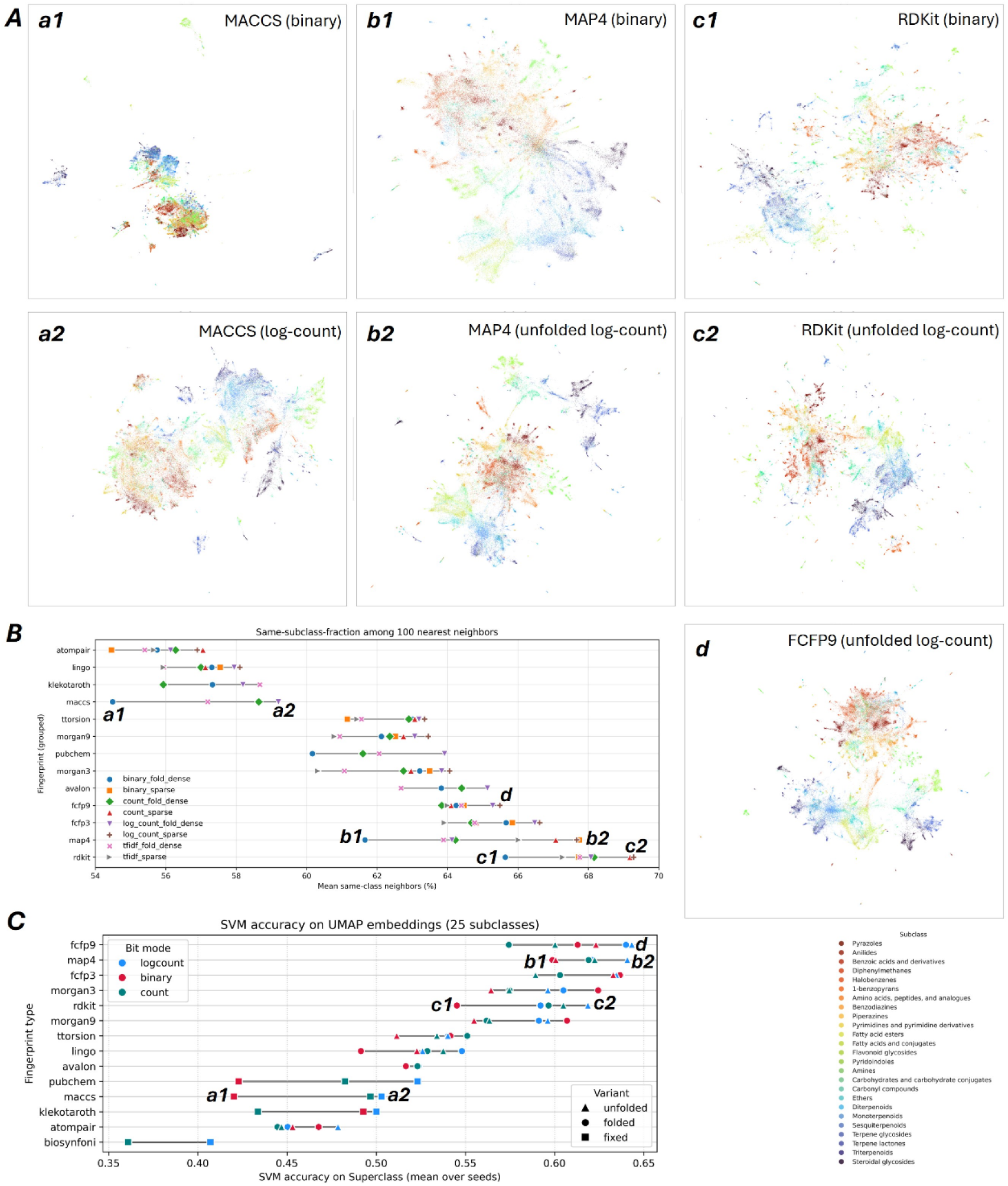
Subclass neighborhood consistency can be one plausible criteria to evaluate a fingerprint’s ability to represent chemically meaningful spaces. (A) Seven examples or UMAP 2D scatter plots for rather low subclass-consistency (MACCS fingerprint) and moderate to high subclass-consistency (MAP4, RDKit, and FCFP9 fingerprints) are shown in the panels a1 to c2 and d. (B) Based on Tanimoto distances a kNN graph was computed for the 25-subclass dataset (75,000 compounds) using pynndescent. The fraction of same-subclass compounds among each datapoint’s k nearest neighbors (here k=100) was computed and the average for each fingerprint type and variant is shown in the bottom part plot. (C) UMAP 2D coordinates where computed based on the Tanimoto-based knn graph (k=100) and SVMs were trained on the 2D coordinates to predict each of the 25 subclasses to give a sense of subclass conherence.

**Figure 12.**
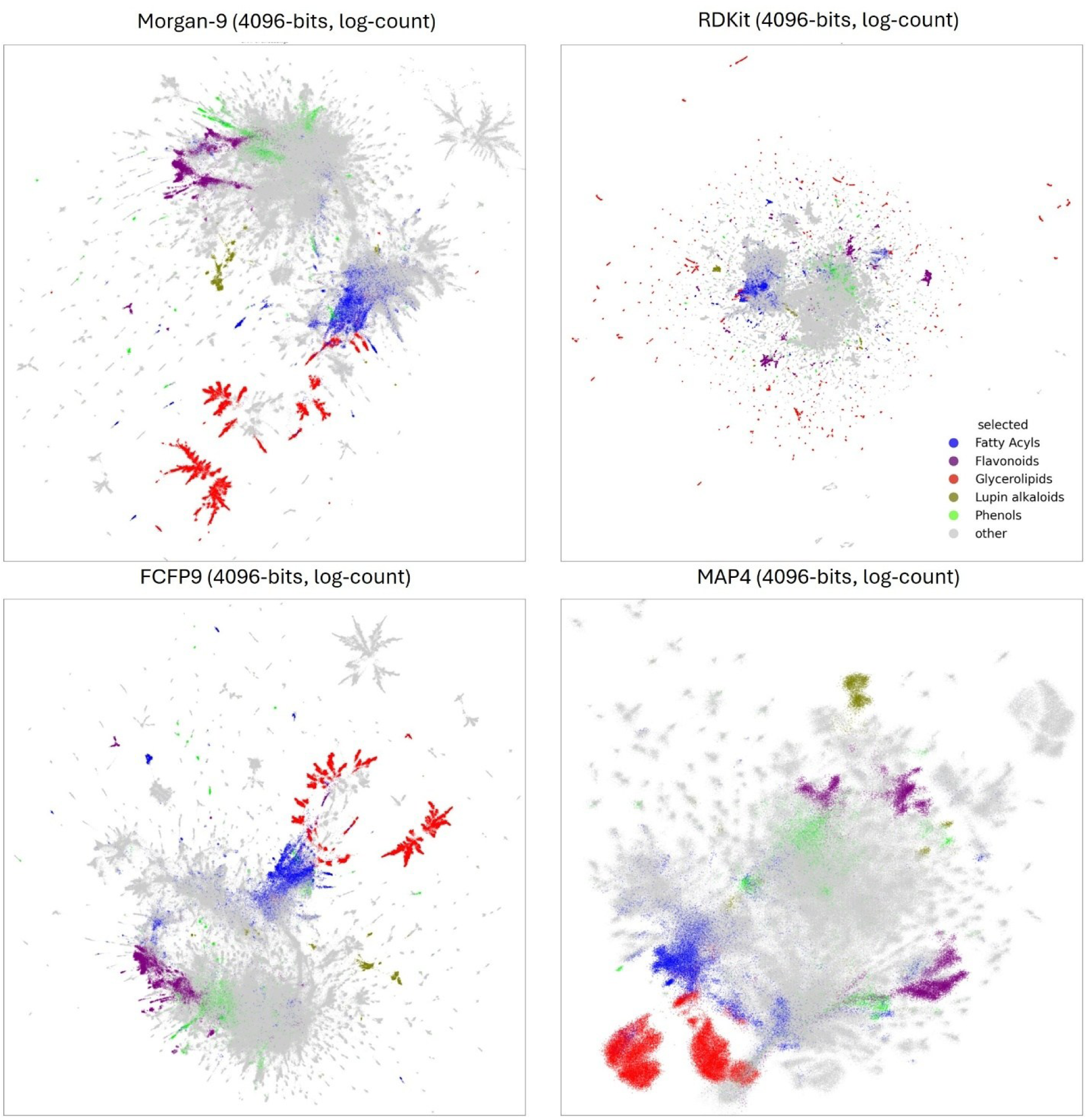
Four examples of UMAP-based chemical space visualizations using the full biostructures dataset. To avoid visual cluttering, only five different chemical classes were selected and colored in the scatter plot. All these UMAP coordinates were computed with chemap using the cuml UMAP implementation which is GPU-optimized. Due to the cuml implementation’s restrictions, folded fingerprints had to be used and distances were computed using cosine (instead of Tanimoto as done in all other parts of this work).

We here tested various variants of each fingerprint type. In addition to the before used binary, count, unfolded binary, and unfolded count variants, we also explored further scaling options: log-count, unfolded log-count as well as TF-IDF scaled vectors to strengthen the effect of less frequent bits in a fingerprint (see methods and supplemental). With the exception of Klekota-Roth, however, the TF-IDF scaling did not improve the subclass neighborhood consistency. The overall best variant choice across fingerprint types was unfolded log-count, or where feasible log-count.

The highest subclass consistency could be measured for RDKit and MAP4 fingerprints. For MAP4, however, we again observed a large discrepancy between folded and unfolded variants with only the latter showing very high subclass consistencies. Follow up consistency values are observed for FCFP (3 and 9) and Avalon fingerprints. Notably lower consistency values are observed for MACCS, Klekota-Roth, LINGO, and Atom Pair fingerprints. In particular Biosynfoni shows generally much lower neighborhood subclass consistency (see supplemental).

When we evaluate the k=10 nearest neighbors (instead of k=100) we see, as expected, that the same-subclass fractions increase. But also, the variant differences shrink, especially the difference between folded and unfolded variants decreases, most notably for MAP4. Still, the overall performance ranking per fingerprint type is only mildly affected with RDKit, FCFP and MAP4 displaying the best overall performance (see supplemental).

The subclass neighborhood consistency further varies considerably among the different subclasses. For some subclasses, compounds consistently remain close together, such as flavanoid glycosides or steroidal glycosides for which we observed typically about 90% of all top-100 neighbors being of the same subclass. Other subclasses such as carbonyl compounds, amines, esters, or monoterpenoids are typically more on the order of 30 to 50 out of 100 nearest neighbors with the same subclass (see more subclass-specific plots in the supplemental).

Finally, we created UMAP^59^-based chemical space visualizations of the 25-subclasses dataset (see methods). Seven of them are shown in Figure 11, which correspond to low and high consistency variants of MACCS, MAP4, and RDKit as well as one example of FCFP9. For MACCS, we see a clear visual improvement with a much more homogeneous distribution of compounds. For MAP4 and RDKit the differences are more subtle, but at least for the case of MAP4 the different subclasses become visibly better separated. This is not unexpected, because UMAP will here also use the k=100 nearest neighbors to compute the position of each compound in 2D, thereby simply reflecting a more consistent neighborhood subclass similarity (for MAP4 this goes from about 62 to about 68% between the two shown variants).

Visual comparison of such scatter plots is often not well suited for judging more subtle differences, yet quantitative evaluations of such visualizations remain inherently difficult and often infeasible. We here applied one technique from Huang et al. ^61^ to at least approximate the substructure coherence in the resulting UMAP 2D coordinates by training support vector machines (SVMs) for predicting the subclasses (see methods, Figure 11 C). This does not exactly follow the neighborhood properties in the original fingerprint space (Figure 11 B), with many general trends between the fingerprints being largely conserved, but a less consistent picture between the different variants. SVMs reached the best accuracies for FCFP, MAP4, Morgan and RDKit fingerprints.

### Large-scale chemical space visualizations

Vector representation, such as molecular fingerprints, as well as the thereby accessible similarity or distance computations, allow the use of modern dimensionality reduction techniques such as UMAP^59^ to create visualizations of large and complex datasets.

We briefly experimented with UMAP alternatives, most notably PACMAP ^67^ which might in some cases provide even slightly better results. Due to computational advantages of UMAP though, especially in a newer GPU-based implementation (cuml)^60^, we kept using UMAP. We provide a simple implementation in chemap, which, given a GPU and enough RAM, allows to compute chemical space visualizations for more than a million compounds in a matter of minutes, with the initial fingerprint computation as main computational bottleneck.

## Discussion

Molecular fingerprint–based similarity measures and several of the limitations we observe are not exactly novelties. However, with fingerprints being used for a growing range of tasks, with growing dataset sizes, the criteria for suitable similarity measures must evolve. Our results provide guidance to avoid common pitfalls and make informed choices regarding fingerprint selection in future studies.

### Conscious fingerprint selection

We would like to start by stating the obvious: different fingerprints will often mean entirely different concepts of what similar molecules will look like, thereby touching on the introduction question of *What does it mean for two molecules to be similar?*

Commonly used fingerprints, such as RDKit or Morgan fingerprints, are often selected arbitrarily ^20^, yet we found their similarity scores behave very differently. We neither can, nor want, to judge which fingerprint type functions best for a particular dataset or research question. But the choice might often be more fundamental than anticipated.

A substantial fraction of pairs that one fingerprint-based score might place in the top 5% may fall below the median for another score (Figure 3). This alone makes the fingerprint choice highly relevant for tasks that depend on the full range of similarity scores, notably when those are used as optimization target in the training of machine-learning models as it is common practice ^8,50,64,68,69^. Similarly, search and ranking results vary substantially with fingerprint choice (Figure 7).

While our study covers several key fingerprint types, it cannot claim to cover the full range of possibilities. Common, established libraries such as RDKit^32^, CDK^70^ scikit-fingerprints ^54^, and chemfp^71^, already provide a large choice of different fingerprint algorithms, often coming with multiple key parameters. Additional variations tested in this work, such as folded, unfolded, or frequency-folded, or bit scaling such as TF-IDF or log counts, will increase the number of combinations even further. We hope that this work will help the cheminformatics community to find suitable benchmarking approaches for future large-scale comparisons.

### Counts your bits

All selected fingerprint types allow count variants. Those will ideally consist of precise integer counts of the respective substructures contained in the fingerprint. Many fingerprints such as all dictionary-based types, but also circular Morgan fingerprints natively support count variants, offering a straightforward switch from presence/absence to occurrence frequency. For other fingerprints, such as the path-based RDKit fingerprint ^32^ or scikit-fingerprints ^54^, these counts, however, do not necessarily reflect true substructure frequencies. Because the algorithm does not de-duplicate overlapping features, structures containing ring systems or internal loops, such as aromatic rings, can yield inflated counts where the same motif is registered multiple times.

Exploring atom or bond disjoint variants would be an interesting future road but was beyond the scope of this work.

Conceptually, count vectors may “overreact” to highly repetitive motifs, for instance, long aliphatic chains in lipid-like molecules. This can, in principle, skew similarity metrics towards compounds that share generic repetitive features rather than chemically distinctive ones. Over all our benchmarking experiments, however, results for count vectors were consistently better, or in a few cases on par, when compared to binary vectors for all tested fingerprint types. Count variants of each fingerprint type in our selections are vastly more specific, as seen in our fingerprint duplication evaluation (Figure 2). They deliver chemical similarities with a smaller gap between small and large molecules (Figure 6). The similarities also align notably better with the graph-based rascalMCES scores (Figure 9). Some prediction tasks, such as the bioactivity prediction we tested, seem to largely depend on only the presence or absence of substructures so that substructures counts do not add much relevant additional value (but also don’t hurt). But for the chemical class prediction and also the subclass neighborhood consistency, we observed often far better performance for count variants (Figure 10 and Figure 11). This also aligns with reports demonstrating improved performance of count-based descriptors in machine learning tasks and environmental screening models ^22^. Unless very specific tasks or particular computational constraints require to use of binary fingerprints variants, we therefore clearly suggest to **default to count variants** even though most fingerprint types in our selection usually come in implementations with the binary variant as standard settings, or are most commonly used in binary mode (such as, for instance, in Boldini et al. ^17^).

### Count scaling

While the benefit of counting substructures is very consistent and notable, further adjustments to those counts cannot be judged so generally. The performed subclass or bioactivity prediction tasks are not suited to assess this aspect. We defaulted to logarithmic scaling for all such models to avoid overly large counts during training. Moreover, given the non-linear nature of the used neural networks, we would also not expect drastic differences between counts and log counts.

Comparing fingerprint-based similarities to rascalMCES (Figure 9), log counts performed worse than count variants for some fingerprint types, and slightly better than count variants for others. For most tasks explored in this work we therefore cannot give any general recommendation.

The only clear exception is subclass neighborhood consistency, where log count variants were either on par or better than other count or binary variants (Figure 11). We therefore expect that tasks such as clustering and dimensionality reduction, e.g., for chemical space visualizations, might benefit from using log counts.

Drawing analogies to common approaches in natural language processing, we further hypothesized that scaling fingerprint bits based on their frequency by TF-IDF might further improve such nearest neighbor-based analyses or visualizations. This, however, could not be confirmed by our latest experiments. With few exceptions, count or log count variants performed better than TF-IDF scaling regarding subclass neighborhood consistency (Figure 11).

### Folded, unfolded, and frequency-folded variants

Unless for dictionary-based fingerprints, bit collisions are a well-known issue with fixed-length fingerprints. But while some studies note considerable decreases in performance ^72^, others find no noteworthy effects^1^. We have the impression that bit collisions are often treated as a relatively minor, hypothetical issue that can be easily mitigated by adhering to common vector sizes.

With growing dataset sizes, and depending on the fingerprint choice, however, our experiments show that bit collisions can lead to highly detrimental effects. In general, this will mostly affect fingerprints with high occupation rates, such as RDKit or MAP4 (Figure 1 and Figure 5).

Common choices for the fingerprint size are 1024 or 2048 bits, which are also the default value in libraries such as RDKit ^32^. We have chosen an even more optimistic baseline of 4096 bits, and yet we observed many arbitrary, unpredictable results and failures. In particular for larger compounds, MAP4 displayed a very high bit occupation and often wrongly assigned pairs of two larger compounds a high chemical similarity (Figure 3) when fixed-sized vectors were used.

A comparison of similarities between 37,811 compounds based on 4096-bit and unfolded fingerprints revealed that for a large fraction of pairs, the classical approach of using a fixed-sized bit vector led to a very considerable overestimation of similarities when using RDKit or MAP4 fingerprints (Figure 4). As a result, when compared to graph-based rascalMCES scores, RDKit and MAP4 fingerprints correlated notably (or, for MAP4, even drastically) better than their folded counterparts (Figure 9). Both fingerprints further also showed much better consistency of scores between small and large molecules (Figure 6). This can be attributed to the higher number of bits with very high bit occupation rates of these two fingerprint types (Figure 1). The more pronounced this effect, the more it will raise the average Tanimoto scores, which is in line with much earlier calculations ^73^ and explains the generally higher Tanimoto scores we observed, but also the higher mass dependency of those fingerprints. For both types, especially for MAP4, we would hence **recommend using unfolded variants** or at least much larger vector sizes (e.g., combined with sparse datatypes) when applied to large, chemically heterogeneous datasets. For fingerprints with lower bit occupation rates, we found that using unfolded implementations even improved similarity computation times while eradicating the risk of potential bit collisions. This performance gain will likely no longer hold when shifting to much larger compounds such as peptides. For high bit occupation fingerprints such as RDKit and MAP4, a sparse implementation can sometimes lead to fingerprints with thousands of bits.

Moving to much larger compounds might then lead to computational challenges. Fingerprints with low bit occupation rates such as LINGO, FCFP-3, or Morgan-3 fingerprints, consistently show only very moderate differences between folded (4096-bit) and unfolded variants (e.g., Figure 6 and Figure 9), representing a low bit collision risk when used in folded variants.

We further hypothesized that bit collisions would be detrimental for downstream tasks such as training machine-learning models, because input values occasionally point to entirely different substructures. For such tasks, using unfolded fingerprints is technically not feasible. To still keep a fixed-sized fingerprint vector, we therefore experimented with a “frequency-folding” by simply only keeping the most occupied bits. This indeed led to notably better performance for Morgan and FCFP fingerprints, but decreasing model accuracies for RDKit fingerprints and only negligible effects for MAP4 or also Atom Pair (Figure 8). We speculate that the very high average bit occupation of RDKit fingerprints, followed by the one of MAP4 and Atom Pair fingerprints (Figure 5) results in a significant information loss when only keeping the most occupied bits (here: 4096) which outweighs or even surpasses the avoidance of bit collisions. Future experiments will have to show if this can in turn be compensated, for instance by using larger vector sizes, or by data-driven clustering of similar substructure bits.

### Dictionary-based vs. substructure searching

There is a fundamental difference between dictionary-based fingerprints that are based on a predefined set of substructures, and substructure searching algorithms that will simply collect (and count) all substructures they can find according to certain algorithms. Dictionary-based types not only have the advantage of having a consistent vector size without any risk of bit collisions, they also can be regarded as much more interpretable (or explainable) since each bit clearly represents a predefined molecular substructure. A key question then obviously is on how to select such substructures, and how many. Already by the number of substructures the four types selected in this work differ widely with 39 substructures in Biosynfoni, 158 for MACCS (166 when binary), 757 for PubChem (881 for binary) to 4860 for Klekota-Roth. While their low dimension makes MACCS and especially Biosynfoni highly interpretable, the broad tasks we here performed on large, heterogeneous datasets also reveal that this comes at the expense of not containing comparable amounts of detailed molecular information in their representations when compared to much larger fingerprint types and variants. PubChem fingerprints, on the other hand, worked surprisingly well for our subclass prediction task and showed high correlation with rascalMCES scores, despite their moderate vector size.

On the bioactivity classification task, but especially in terms of compound specificity as measures by fingerprint duplicates, however, all selected dictionary-based fingerprints performed comparatively poorly. This could be rooted in the respective substructure selections, which were typically chosen with respect to particular tasks, or chemical classes or domains ^16,28–30^ whereas we here used larger, chemically heterogeneous datasets (see methods). More substructure, specific to the present compounds, are likely explains the often superior results of fingerprints such as FCFP or Morgan, both frequently even improving for larger substructure sizes.

Nevertheless, in light of the surprisingly large discrepancy between fingerprints with few dozens to fingerprints with thousands of substructure bits, we imagine that re-iterations to design of novel dictionary-based fingerprints might be worthwhile for obtaining task-specific compact yet performant and explainable representations.

### General-purpose fingerprints for broad chemical space representation

From the beginning, and strongly underlined by our various experiments, it seems clear that there will not be one single best choice in terms of fingerprint type and variant. However, when the focus lies on a broad representation of heterogeneous chemical spaces (still limited to small molecules), several candidates performed reasonably well across many of our benchmarking tasks. This includes RDKit and MAP4 fingerprints (when used unfolded), as well as Morgan and FCFP fingerprints, especially in combination with larger radii. A conceptual benefit of Morgan and FCFP fingerprints is that they require far fewer bits, on average about an order of magnitude fewer than RDKit, and about five times fewer than MAP4, making them more memory efficient and often considerably faster for similarity computations.

Those fingerprints are also well-suited for dimensionality reduction-based chemical space visualization using algorithms such as UMAP. We provide two different UMAP implementations in the library chemap to compute Tanimoto-based visualizations for up to a few hundred thousand compounds (CPU based), or a GPU-based versions relying on cosine instead of Tanimoto that can more easily handle very large datasets with more than a million different compounds.

Importantly, the magnitude of effects discussed throughout this work (e.g., folding-induced bit-collision artifacts, compound-size dependence, and fingerprint duplication) depends on the chemical composition of the dataset, including its size distribution and structural diversity. Our datasets were chosen to stress-test fingerprint behavior on broad, biologically relevant and chemically heterogeneous collections (metabolomics/natural products), and the same effects may manifest differently in more constrained libraries such as typical drug-like screening sets. A dedicated evaluation on such collections is therefore a natural extension of this work.

In summary, our results argue for a more deliberate fingerprint selection, which will severely change similarity behavior and downstream results. Across most settings we tested, count fingerprints were preferable to binary variants, and log-scaling was a robust default for neighborhood-based analyses such as chemical space visualization. For fingerprints with high bit occupation, switching to unfolded/sparse variants is advantageous for RDKit and effectively required for MAP4 on broad, heterogeneous datasets.

Finally, our results show that it can be very promising to re-evaluate established defaults: while Morgan-2/3 are commonly used, larger radii and count (or log-count) representations often worked better, and Morgan-9 or FCFP9 fingerprints generally performed well across tasks.

These variants are certainly not so good as to rule them all, but they provide robust starting points for broad chemical space representation.

## Availability of data and materials

Both datasets used in this work are publicly available at https://zenodo.org/records/18682051 All code required to reproduce the presented results can be found on GitHub: https://github.com/florian-huber/molecular_fingerprint_comparisons.

## Supporting information

Huber_Pollmann_count_your_bits_supplemental_v3

## Acknowledgements

We want to thank Andrew Dalke, Niek de Jonge and Justin J.J. van der Hooft for their very helpful feedback on an earlier version of this manuscript.

This work was funded by the Deutsche Forschungsgemeinschaft (DFG, German Research Foundation) – 528775510.

## Authors’ contributions

**FH** conceived the study, designed the experiments, and supervised the project. **FH** and **JP** wrote the code, performed the experiments, analyzed the data, and wrote the manuscript.

