## Supplementary material for "Count your bits: fingerprint benchmarking to assess broad chemical space representation": Huber_Pollmann_count_your_bits_supplemental_v3

#### Fingerprint bit occupation when compared to more compressed 1024-bit variants

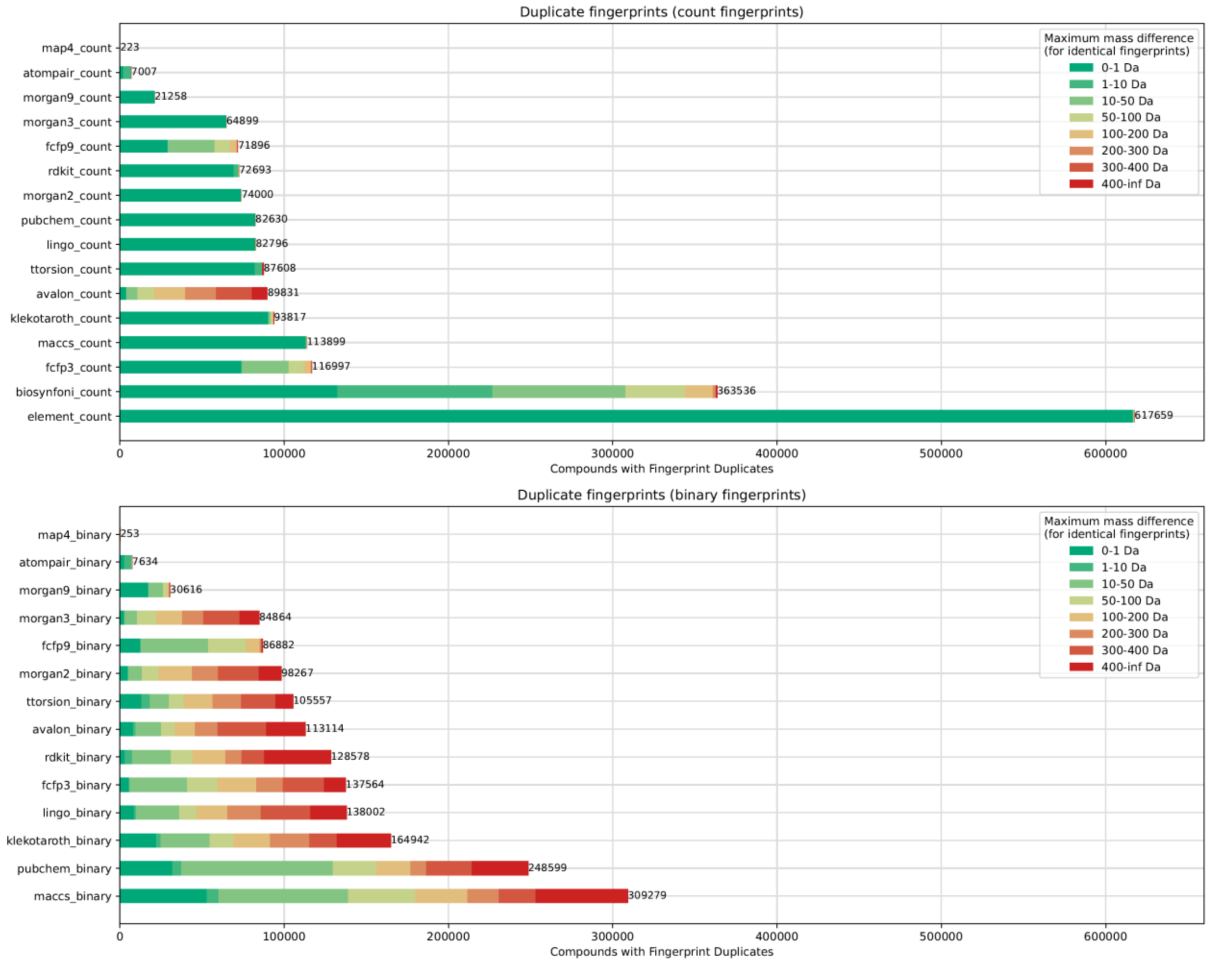


*Figure 1 For different molecular fingerprints, all duplicates in a set of 718,067 unique compounds (biomolecular structures dataset) were counted. For each duplicate fingerprint, we computed the maximum mass difference among compounds sharing that fingerprint which is colored along different bins ranging from 0-1Da mass difference up to mass differences >= 400 Da. Here, unlike in the main article, we also included Biosynfoni, and, as a baseline, a simple element count vector.*

### Examples of high mass difference fingerprint duplicates (MACCS)

mass difference: 947.151 Da

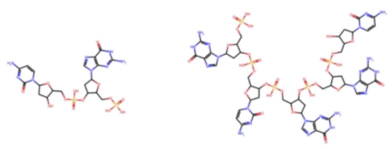

mass difference: 304.116 Da

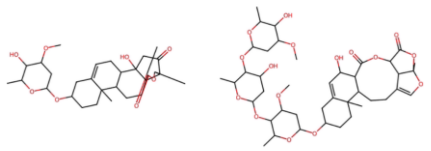

mass difference: 349.978 Da

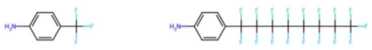

mass difference: 520.595 Da

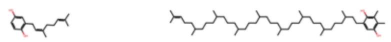

mass difference: 392.438 Da

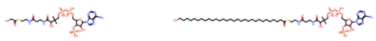

mass difference: 311.642 Da

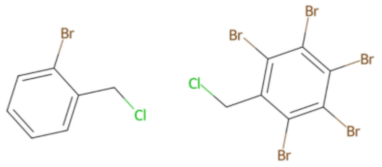


mass difference: 448.501 Da

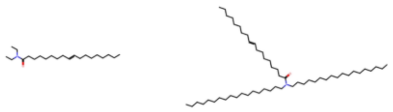

mass difference: 762.271 Da

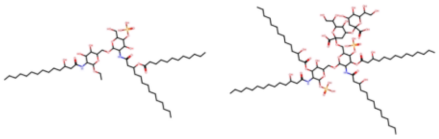

mass difference: 896.301 Da

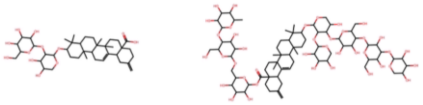

mass difference: 610.248 Da

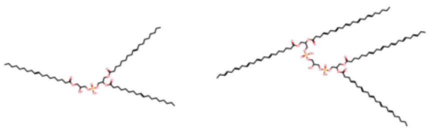

mass difference: 228.093 Da

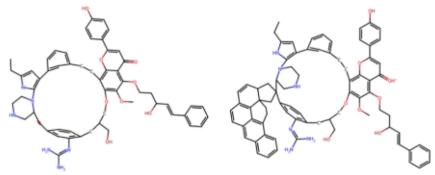

mass difference: 206.203 Da

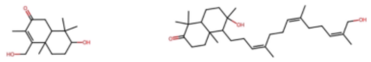

mass difference: 398.242 Da

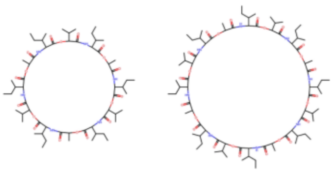

mass difference: 220.219 Da

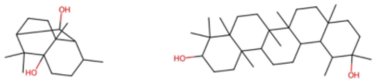


mass difference: 222.235 Da

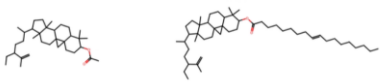

mass difference: 997.127 Da

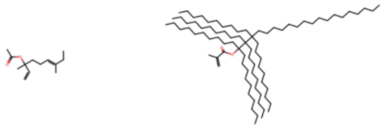

#### Examples of high mass difference duplicates - RDKit binary (4096 bits)

mass difference: 276.245 Da

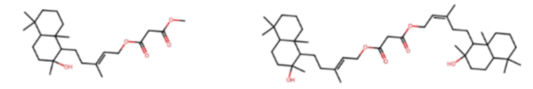

mass difference: 402.131 Da

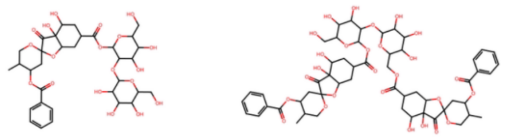

mass difference: 309.205 Da

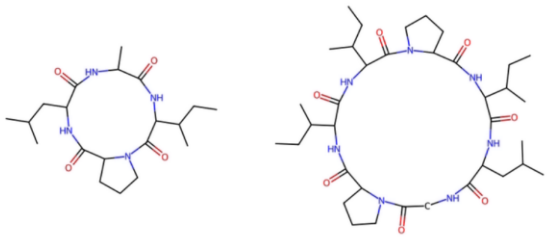

mass difference: 476.459 Da

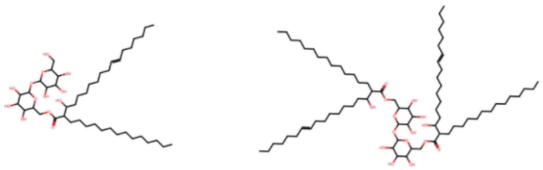

mass difference: 348.266 Da

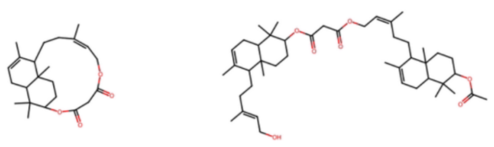

mass difference: 320.111 Da

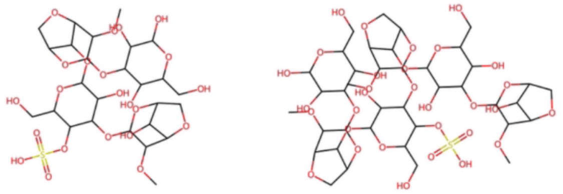

mass difference: 388.043 Da

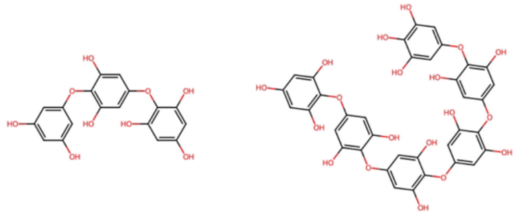

mass difference: 403.937 Da

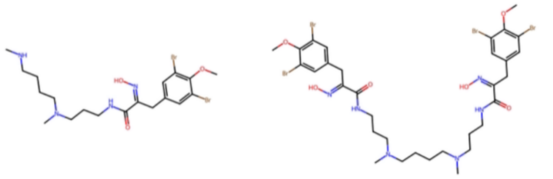

mass difference: 734.673 Da

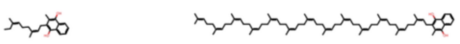

mass difference: 280.313 Da

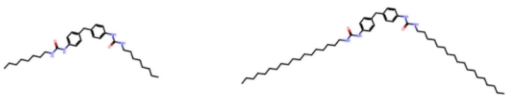

mass difference: 466.124 Da

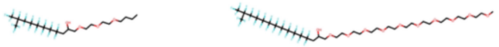

mass difference: 252.282 Da

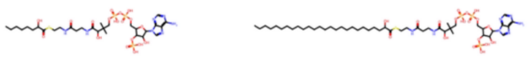

mass difference: 344.344 Da

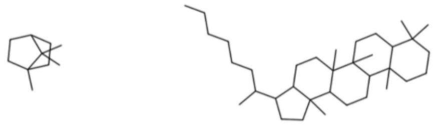

mass difference: 1037.158 Da

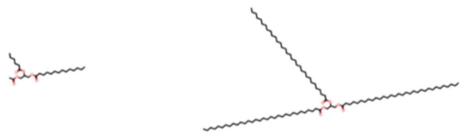

### Mass-sensitivity of different similarity measures

In the main article we used a percentile scaling to better compare the different similarity scores. Here, just for comparison, we also display the overall similarity scores for all compounds <300Da and >500Da in the ms2structures dataset without the percentile scaling.

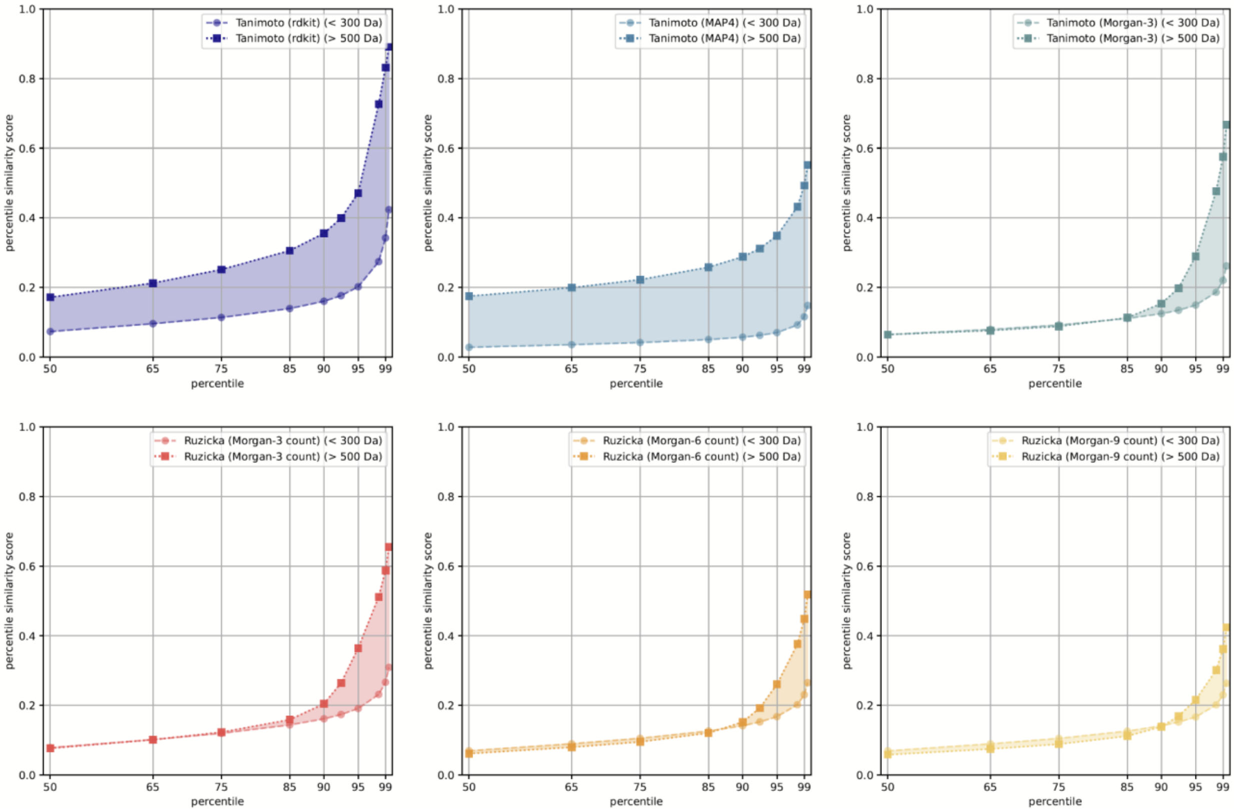


*Figure 2 Fingerprint-based similarities are computed for about 36 million unique pairs between small compounds (< 300 Da) or between larger compounds (> 500 Da). The similarity scores of increasing percentiles are plotted for both the small and larger molecules ranging from 50% up to 99.5%. This is done for Tanimoto scores of rdkit fingerprints ( top left), Tanimoto scores of MAP4 fingerprints (top, middle), Tanimoto scores of Morgan-3 binary fingerprints (top right) as well as Ruzicka scores of Morgan-3 to Morgan-9 count vectors (bottom row). All used fingerprints had 4096 bits.*

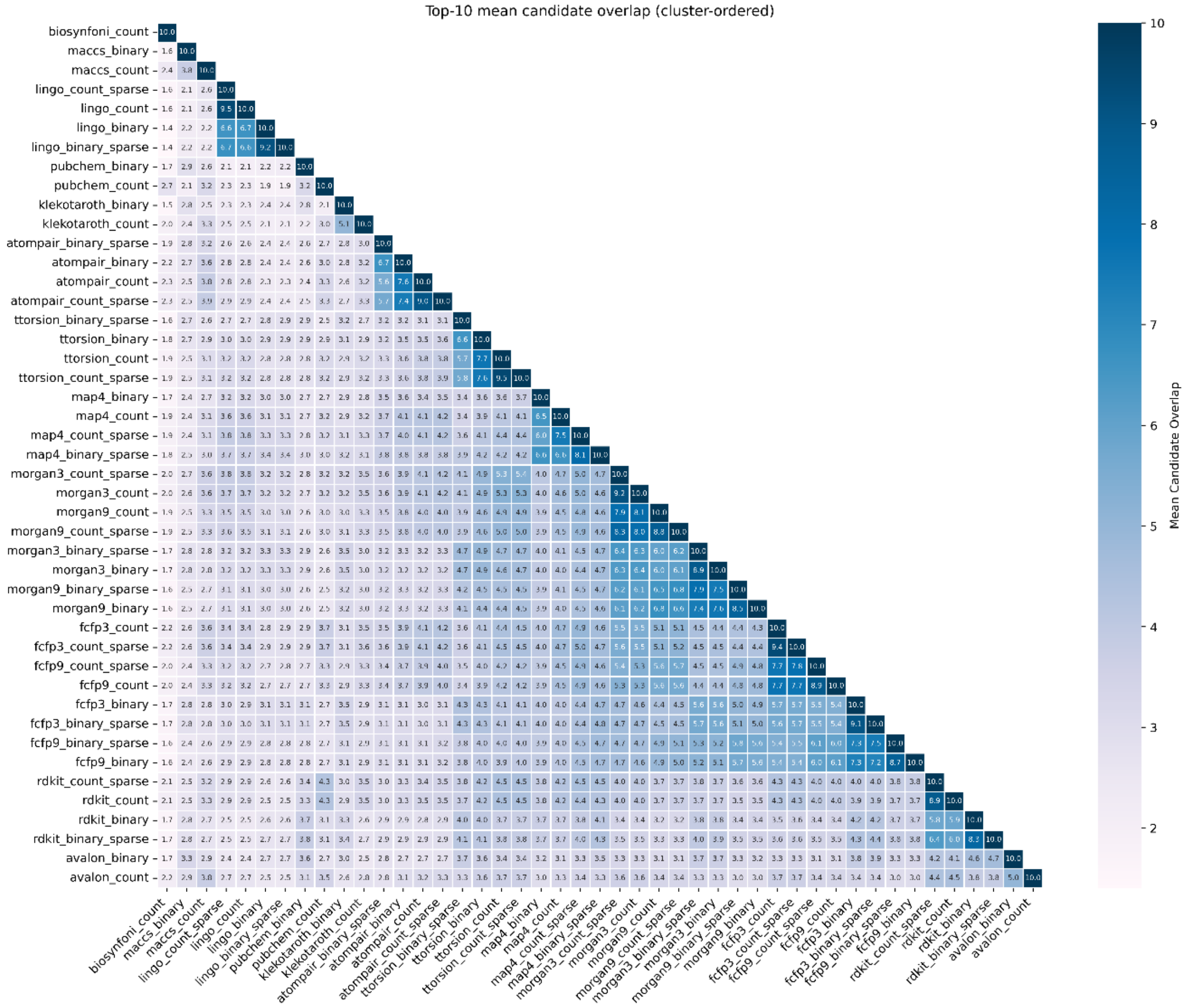


*Figure 3 Heatmap to display all top-10 ranking overlaps between the different fingerprint types and variants. Those values were used to later compute a minimum spanning tree (see main article).*

#### Activity prediction vs. virtual screening experiment

A frequently cited, and adapted, fingerprint evaluation method is the one by Riniker and Landrum [1].

In the context of our attempt to shed a rather broad perspective on fingerprint similarity, we contend that this ranking-based aggregation might introduce two potential issues, one with the aggregation of ranks and one with the underlying screening task. Aggregation of ranks is a commonly applied technique, mostly because it focuses on relative performance per query, is robust to metric outliers, and avoids unjust bias towards methods optimized on high-variance subsets [2]. But converting continuous performance differences into ordinal ranks can obscure the true magnitude of those differences. Even minimal differences may result in a rank change of one, while substantial differences may be treated equivalently if they occupy consecutive rank positions. And, the averaged ranks are not independent quantities but depend on the selection of competing scores.
Secondly, the ranking tasks in the mentioned studies [1], [3], [4] are geared towards detecting compounds with assigned biological activity within a large pool of decoys, thereby simulating

the detection of active compounds. This, however, is not directly related to the conceptual design of most fingerprints, which are rather broad descriptors of overall molecular substructures.
We ran the ranking script from Orsi and Reymond [4] with various sparse Morgan-based fingerprints. Overall ranking performance on the datasets provided for small molecules remained best for the MAP4 fingerprints (see Figure S2 in supplemental material).
Since those mean ranking plots depend on the selection of scores and hence do not allow for a flexible, general benchmarking over many different scores and future adaptations, we here also compare several other performance metrics as used in [4] (Figure 3). Here, too, MAP4 frequently received the best average metrics, though not on all provided metrics.
Interestingly, we found that larger radius Morgan fingerprints, such as sparse log-count Morgan-9 fingerprints perform comparably well on the peptide task, being roughly on par with MAP4 for the mean rank (Figure 4) as well as the difference mean metrics (Figure 6).

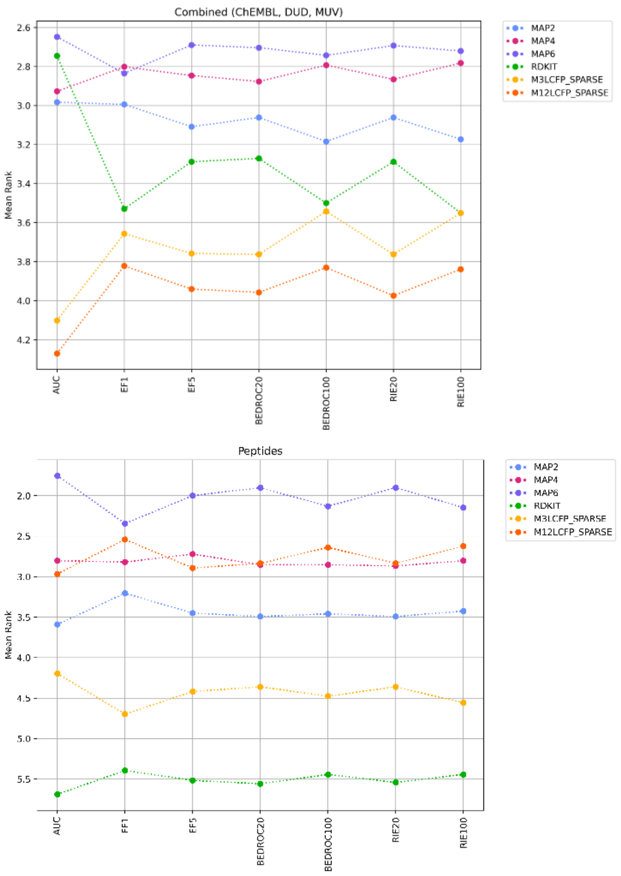


*Figure 4 Mean ranks of fingerprints across all virtual screening datasets as provided in Orsi and Raymond (2024). Left: For the three datasets for small molecules (ChEMBL, DUD, MUV). Right: For the peptides datasets.*

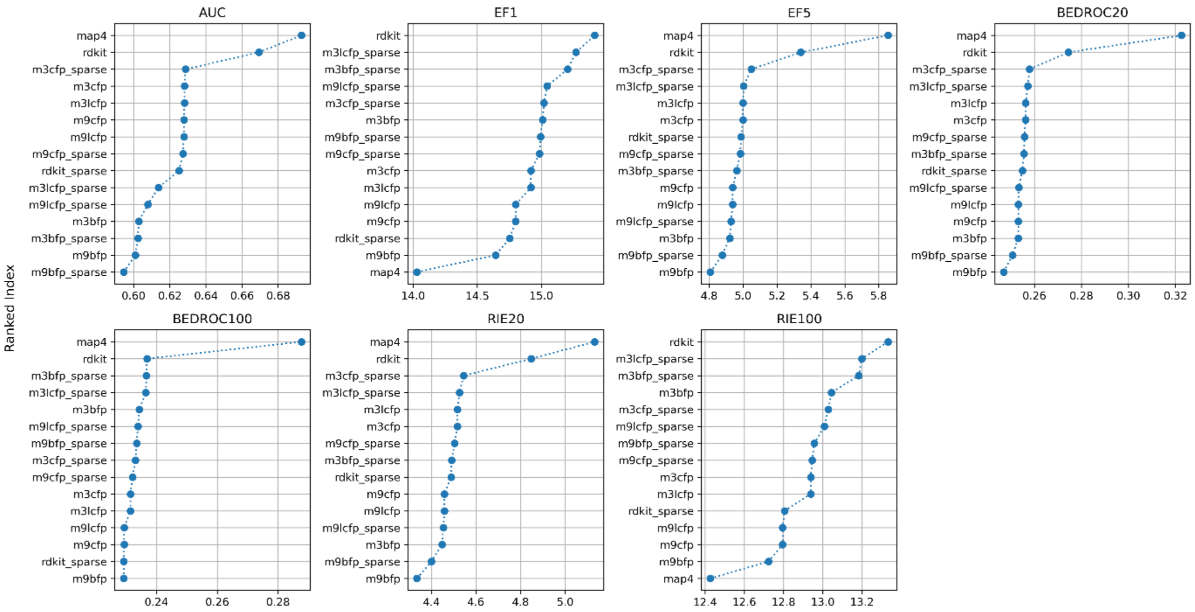


*Figure 5 Mean scores of fingerprints across the 3 virtual screening datasets for* ***small molecules (ChEMBL, DUD, MUV)*** *as provided in Orsi and Raymond (2024). For each subplot the respective values were ordered in descending order to allow identifying which fingerprints performed well or poorly according to various metrics.*

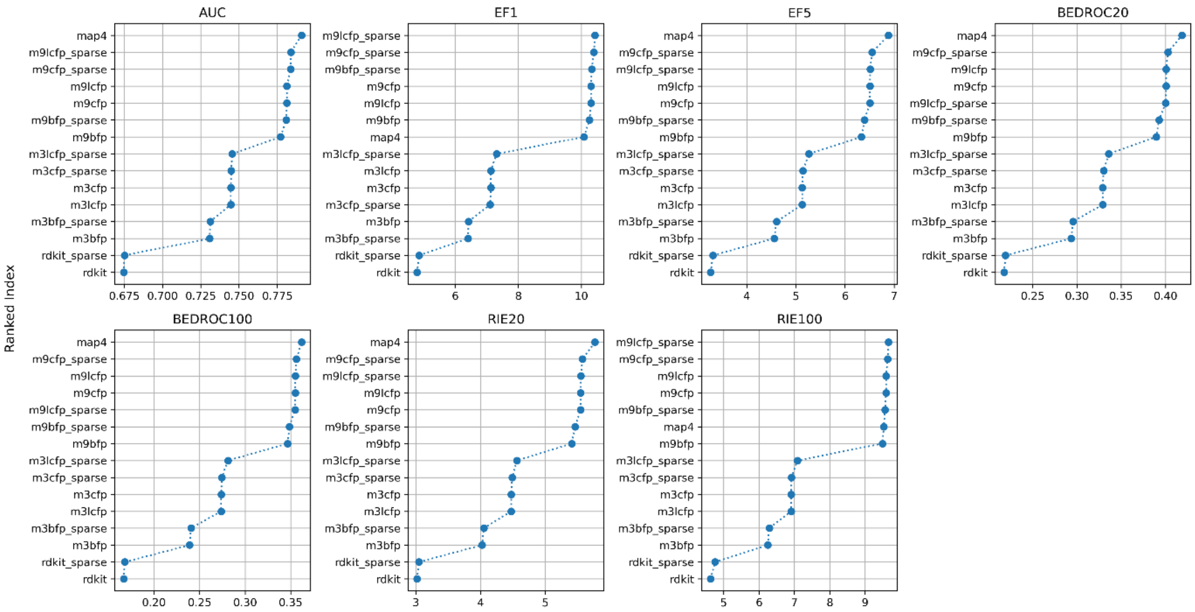


*Figure 6 Mean scores of fingerprints across the 2 virtual screening datasets for* ***peptides*** *as provided in Orsi and Raymond (2024). For each subplot the respective values were ordered in descending order to allow identifying which fingerprints performed well or poorly according to various metrics.*

#### Subclass prediction task (120-subclasses dataset)

In the main article, we did not include the results for Biosynfoni for the 120 subclass prediction task because it became clear, that the fingerprint was not providing enough information for the models to compete with the other, much larger, fingerprint types. We expect, that it might still perform much better on a subset of the subclasses we trained on, but we have not done any steps to evaluate this any further.

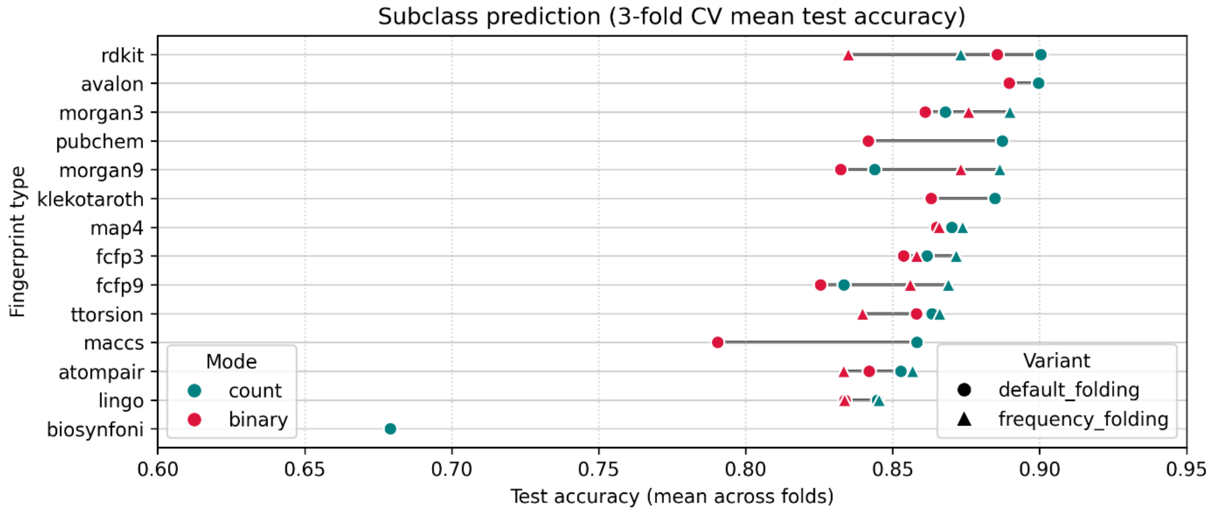


*Figure 7 In the main article we excluded Biosynfoni because it could not compete with the other, larger, fingerprint types and variants.*

As mentioned in the main article, our goal was not to train the most performant models possible. We used, on purpose, only a subset of the available data. And we also did not do very excessive parameter scans. So we expect that the actual accuracy ratings can be improved further by using more training data and spending more time to optimize model architecture and training procedure.
As a sanity check, however, we did train a much simpler architecture with only one hidden layer (3000 nodes). While some accuracy values changed slightly, the overall results were largely on par with the deeper model used in the main article.

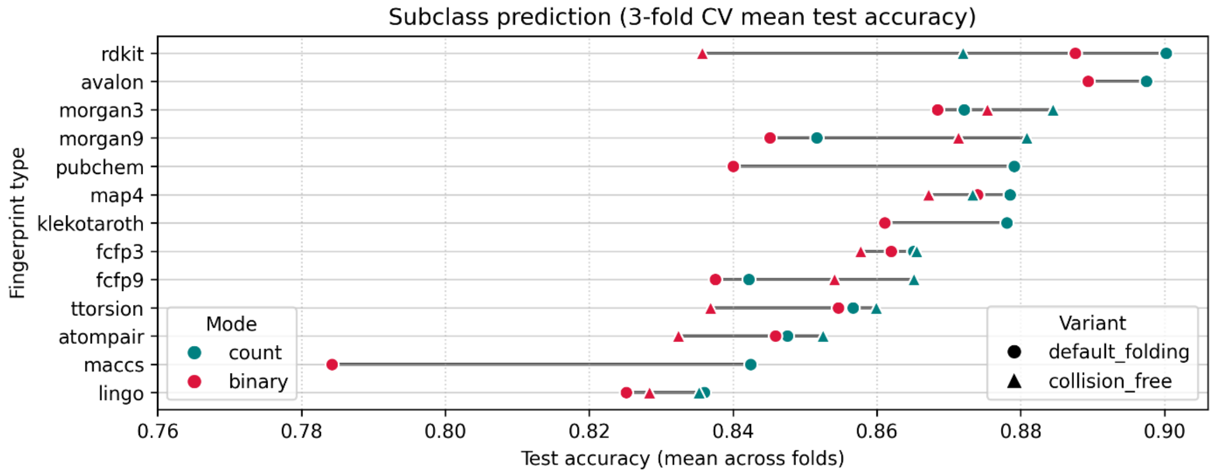

#### Subclass neighborhood consistency (25-subclasses dataset)

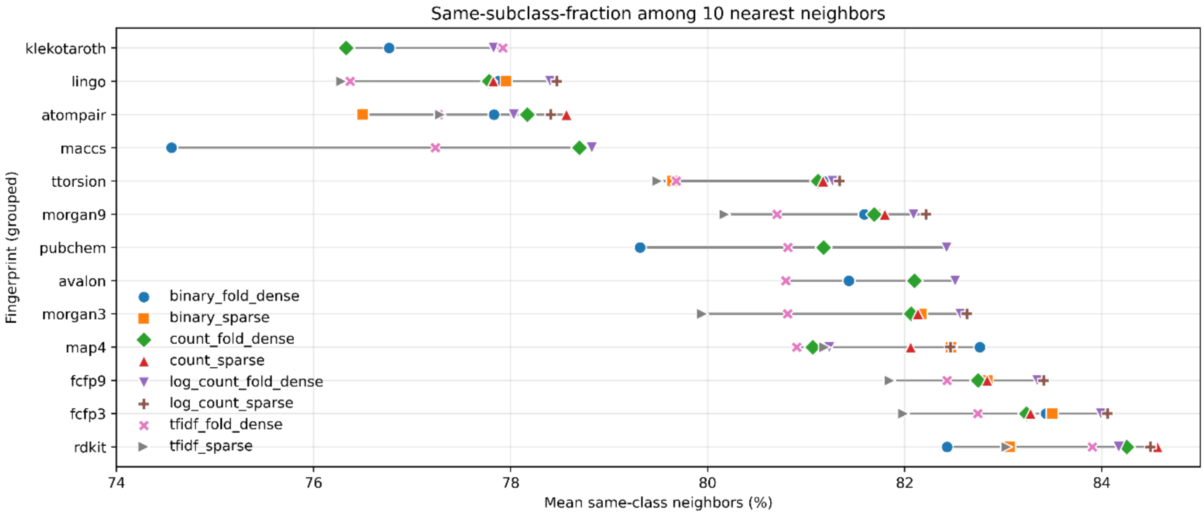


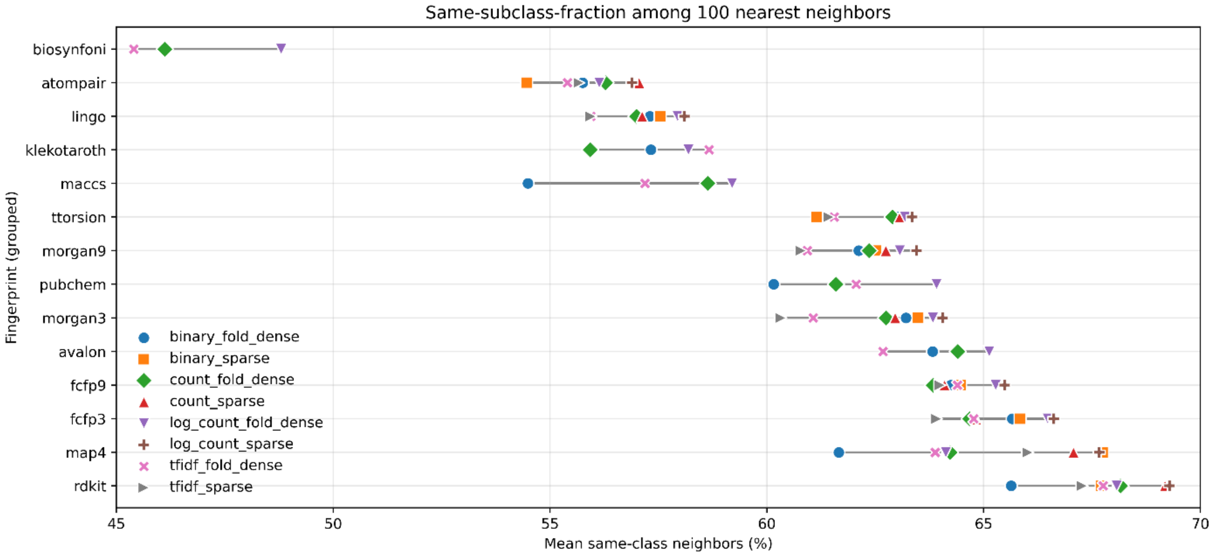


*Figure 8 Plot of all fingerprint types, including Biosynfoni on the subclass consistency.*

The following plot show the measured same-class consistency for each of the 25 selected subclasses.

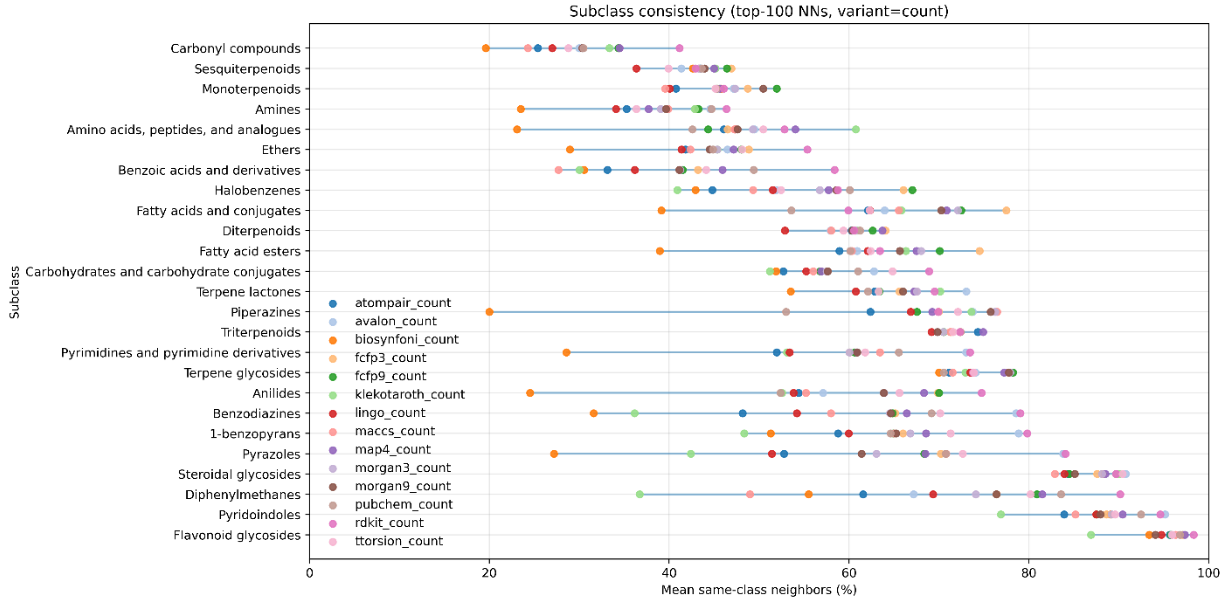

Subclass consistency (top-100 NNs, variant=count)
Carbonyl compounds
Sesquiterpenoids
Monoterpenoids
Amines
Amino acids, peptides, and analogues
Ethers
Benzoic acids and derivatives
Halobenzenes
Fatty acids and conjugates
Diterpenoids
Fatty acid esters
Carbohydrates and carbohydrate conjugates
Terpene lactones
Piperazines
Triterpenoids
Pyrimidines and pyrimidine derivatives
Terpene glycosides
Anilides
Benzodiazines
1-benzopyrans
Pyrazoles
Steroidal glycosides
Diphenylmethanes
Pyridoindoles
Flavonoid glycosides
Subclass
atompair_count
avalon_count
biosynfoni_count
fcfp3_count
fcfp9_count
klekotaroth_count
lingo_count
maccs_count
map4_count
morgan3_count
morgan9_count
pubchem_count
rdkit_count
ttorsion_count
0
20
40
60
80
100
Mean same-class neighbors (%)


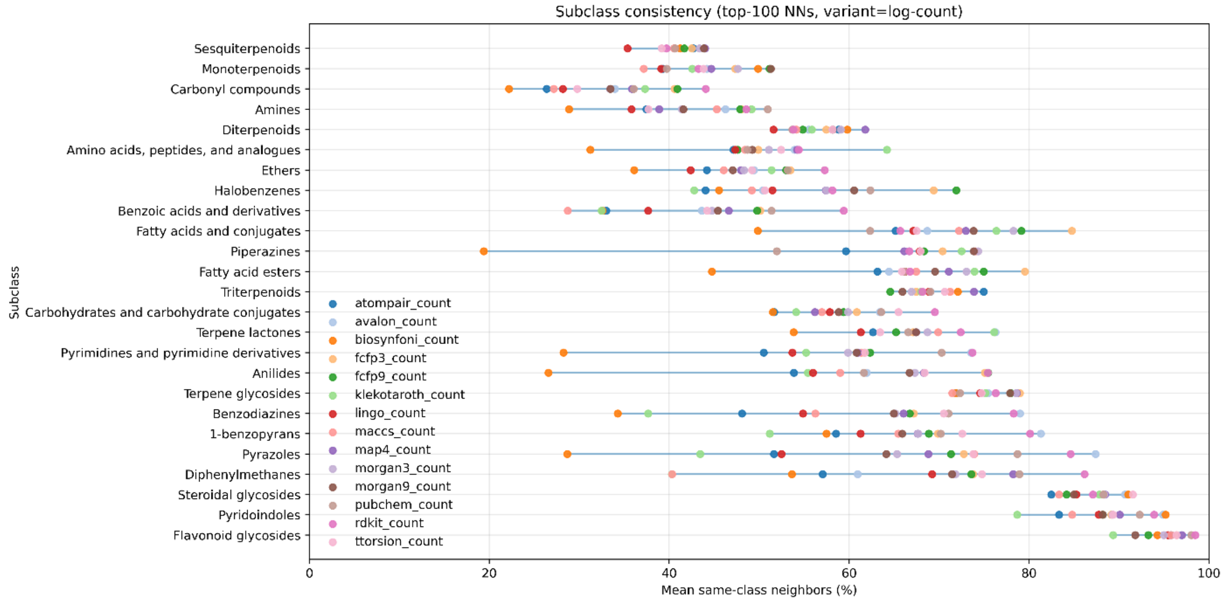

Subclass consistency (top-100 NNs, variant=log-count)
Sesquiterpenoids
Monoterpenoids
Carbonyl compounds
Amines
Diterpenoids
Amino acids, peptides, and analogues
Ethers
Halobenzenes
Benzoic acids and derivatives
Fatty acids and conjugates
Piperazines
Fatty acid esters
Triterpenoids
Carbohydrates and carbohydrate conjugates
Terpene lactones
Pyrimidines and pyrimidine derivatives
Anilides
Terpene glycosides
Benzodiazines
1-benzopyrans
Pyrazoles
Diphenylmethanes
Steroidal glycosides
Pyridoindoles
Flavonoid glycosides
Subclass
atompair_count
avalon_count
biosynfoni_count
fcfp3_count
fcfp9_count
klekotaroth_count
lingo_count
maccs_count
map4_count
morgan3_count
morgan9_count
pubchem_count
rdkit_count
ttorsion_count
0
20
40
60
80
100
Mean same-class neighbors (%)

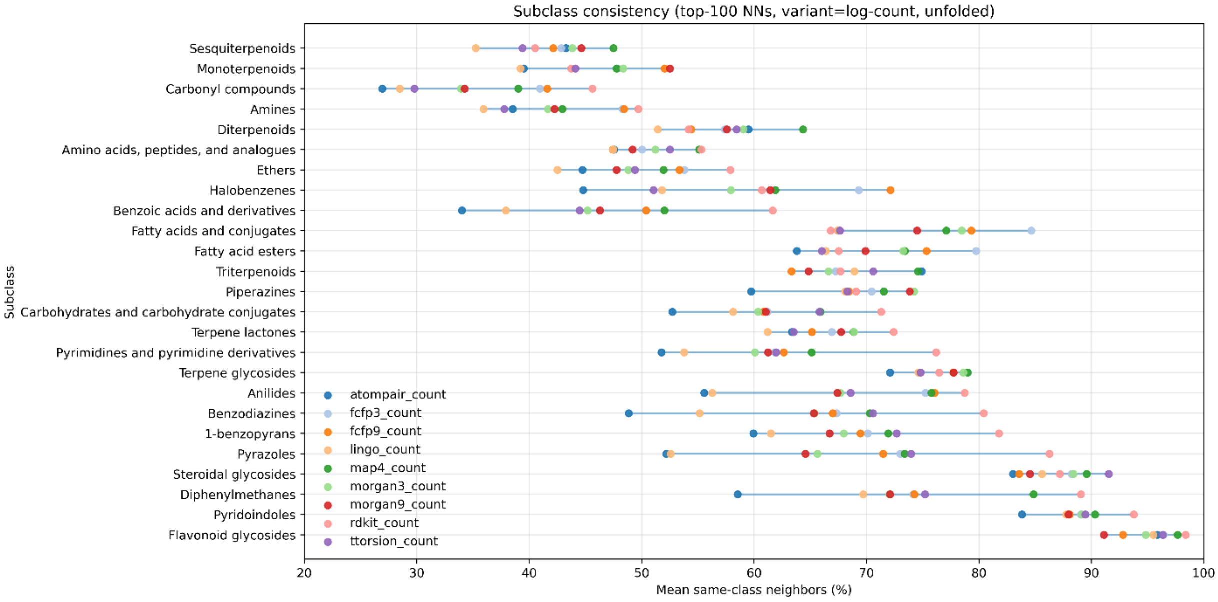


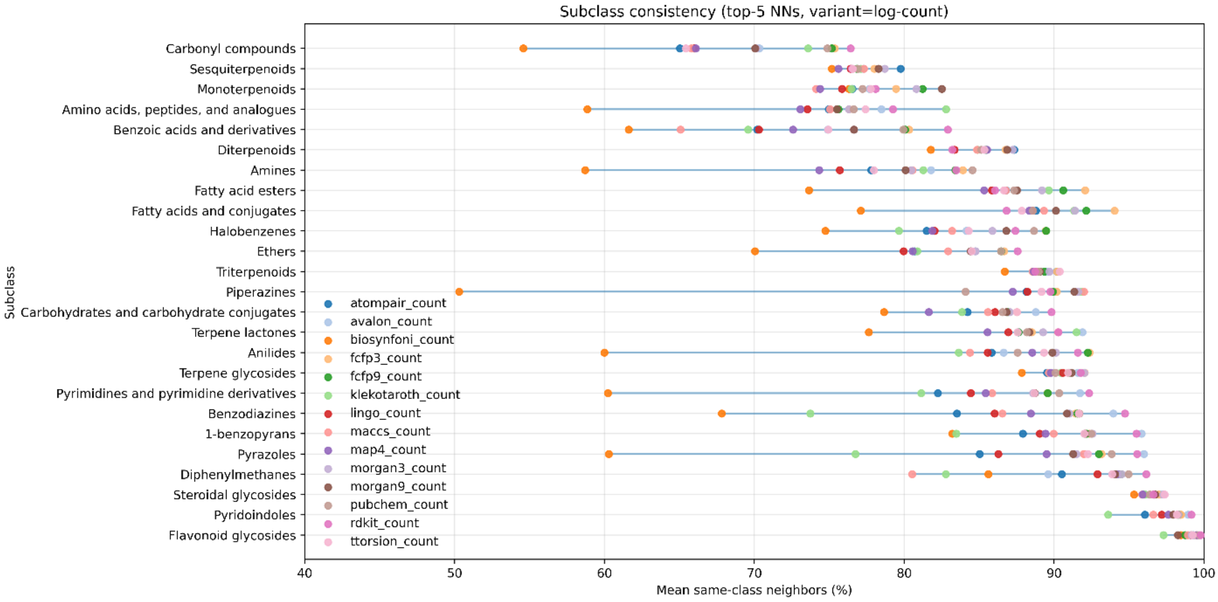


#### Exploring individual bit counts and weighing options

The bit occupation distributions in shown in the main article (**Error! Reference source not found.**) displays that some fingerprint bits occur much more often than others. In the logic of both binary and count-based fingerprints, such as the RDKit and Morgan fingerprints, but also MAP4, all fingerprint bits are equally important. This means that for a similarity computation, the presence or absence of a very common substructure counts as much as the presence or absence of a rarely occurring substructure.

We weighted each fingerprint bit using the inverse document frequency (IDF, see Methods). This was done using sparse RDKit and Morgan fingerprints to avoid bit collisions. As expected, the most frequent substructures for Morgan fingerprints were of radius 0 or 1 and exclusively contained the atoms H, C, O, N (Figure 9). The most frequent substructure, an oxygen atom with a double bond, was found in about 84% of all fingerprints in the ms2structures dataset and in about 81% of all fingerprints in the biostructures dataset (Figure 10). RDKit fingerprints contain many more substructures that occur in a large fraction of all molecules, in agreement with **Error! Reference source not found.** (main article). The most frequent substructures here are C-C sequences of various lengths as well as other trivial motifs like C-O or C=O (Figure 11).

On both the ms2structures and the biostructures dataset, we computed sparse binary Morgan-3 fingerprints for all compounds. The total number of occurrences of each bit (=substructure) across the respective dataset was then taken to compute the IDF weights for each bit.

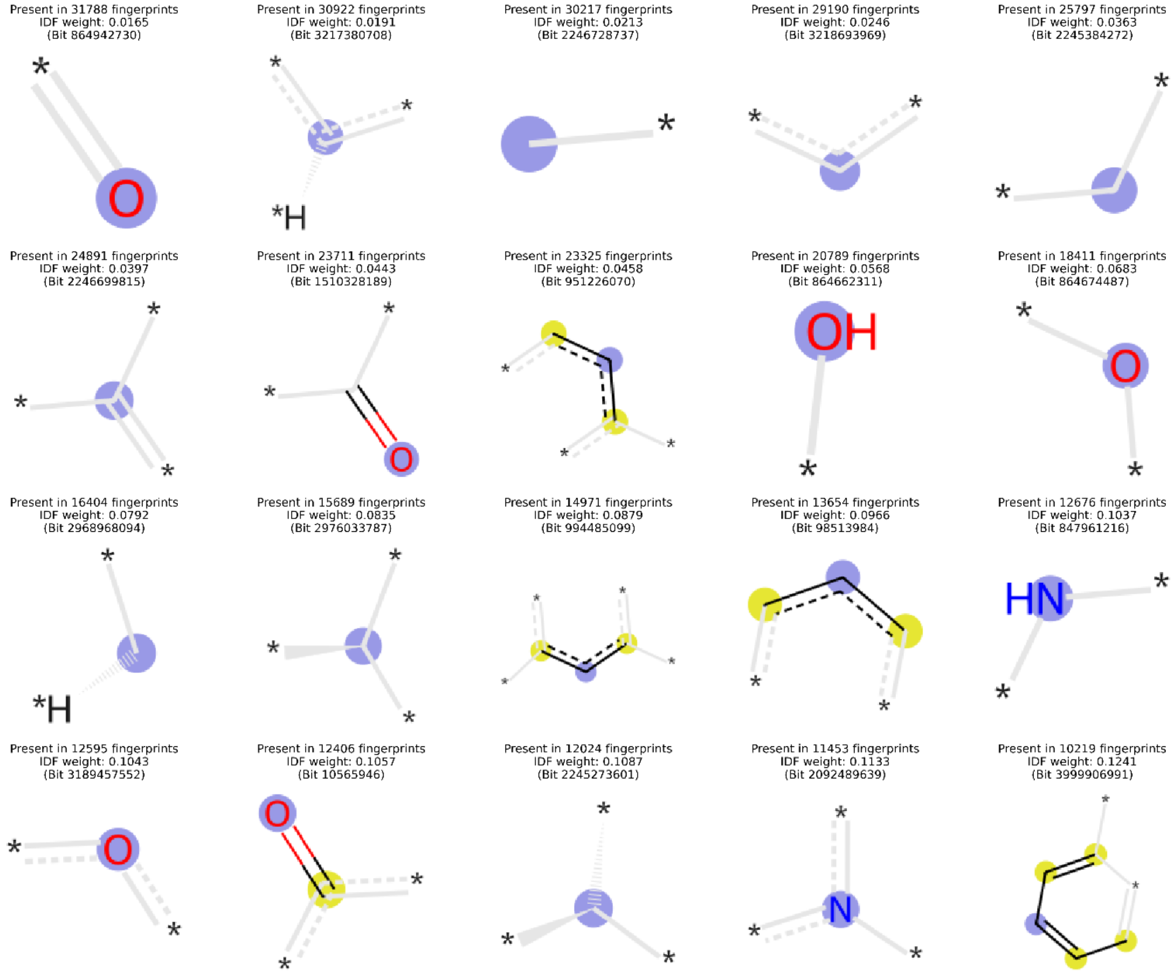


*Figure 9. The 20 most frequently found Morgan-3 fingerprint bits in the **ms2structures dataset**. To avoid bit collisions, sparse Morgan-3 fingerprints were used and bit occurrences were counted. The blue circle marks the center atom.*

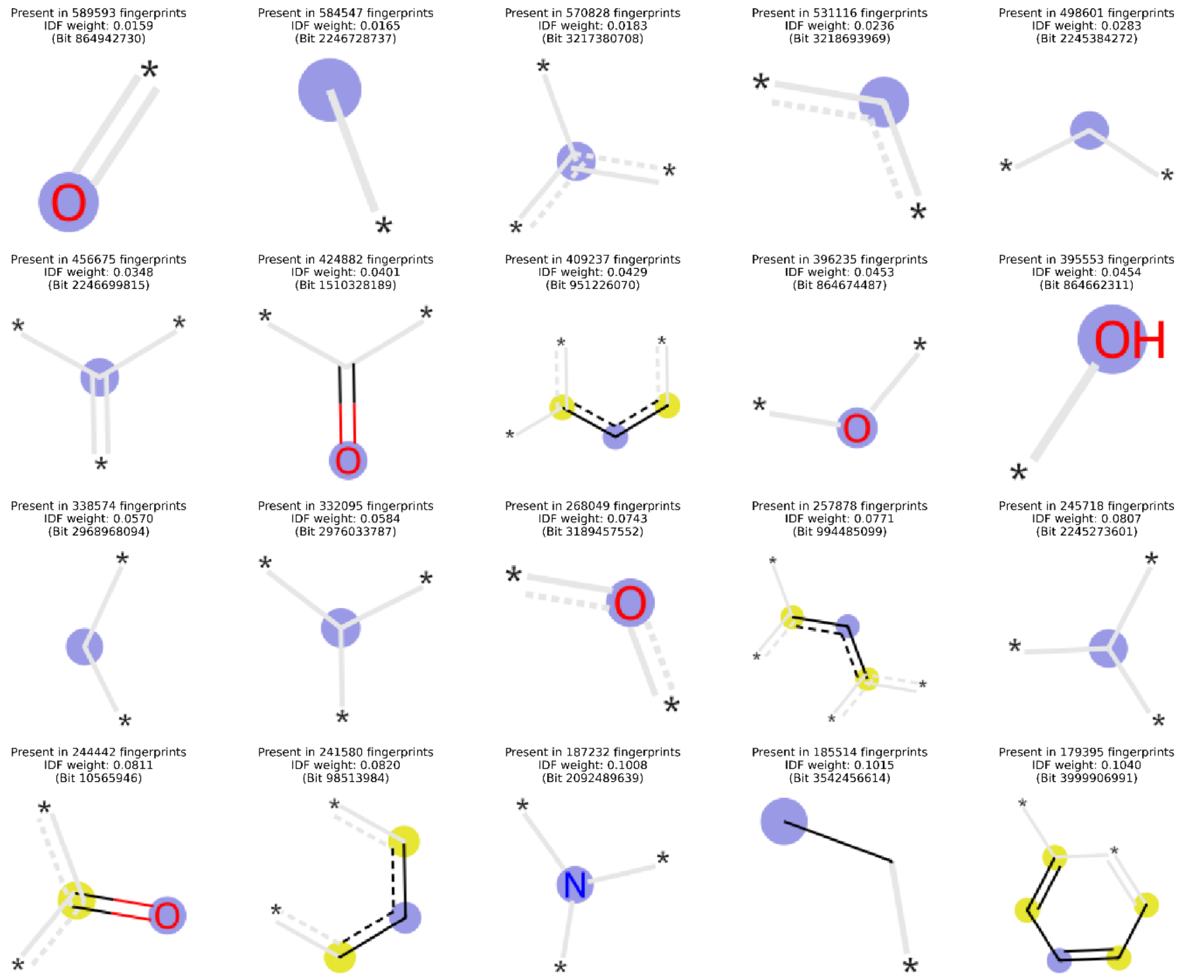


*Figure 10 The 20 most frequently found Morgan-3 fingerprint bits in the* ***biostructures dataset****. To avoid bit collisions, sparse Morgan-3 fingerprints were used and bit occurrences were counted. The blue circle marks the center atom.*

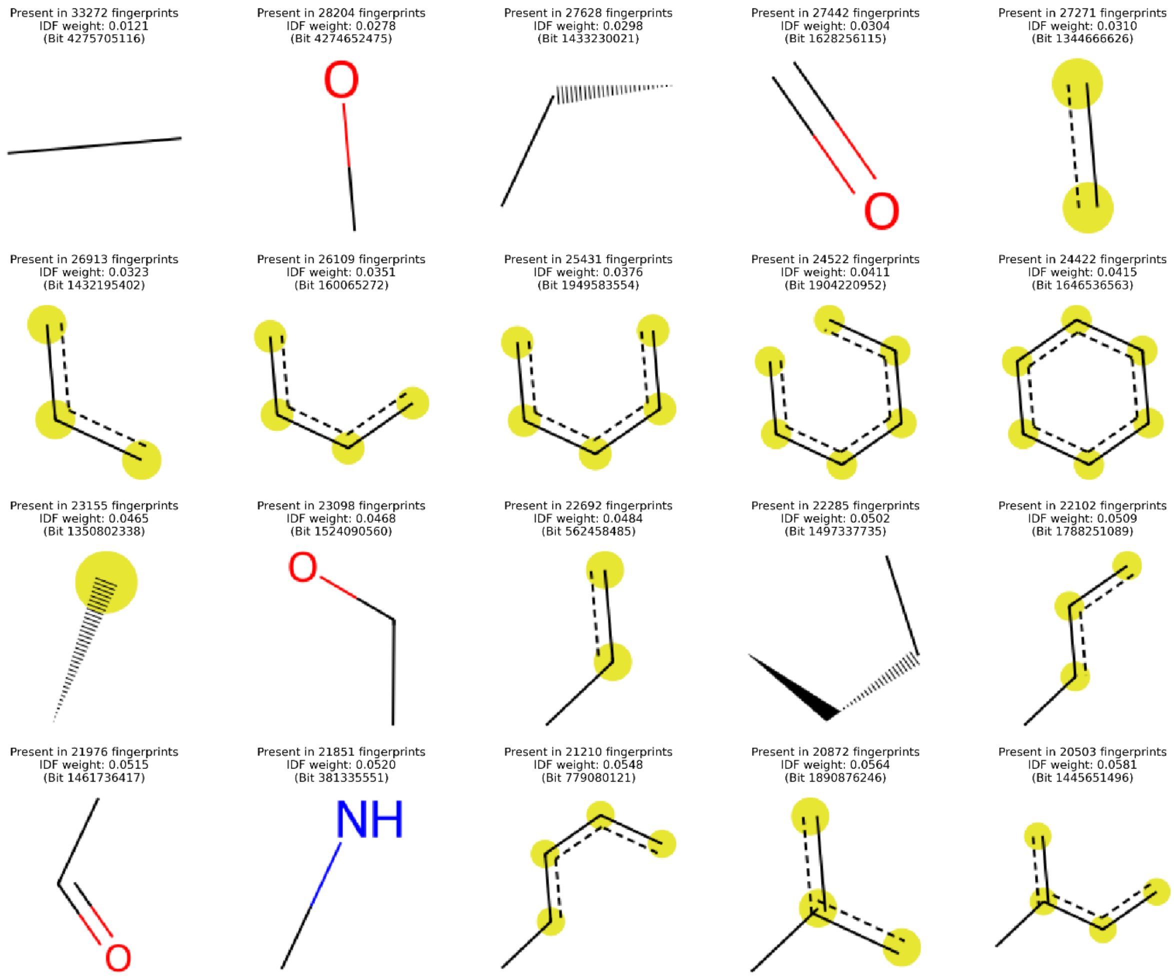


*Figure 11. The 20 most frequently found RDKit fingerprint bits in the **ms2structures dataset**. To avoid bit collisions, sparse RDKit fingerprints were used and bit occurrences were counted. When yellow circles are used, they represent the atoms belonging to the respective paths.*

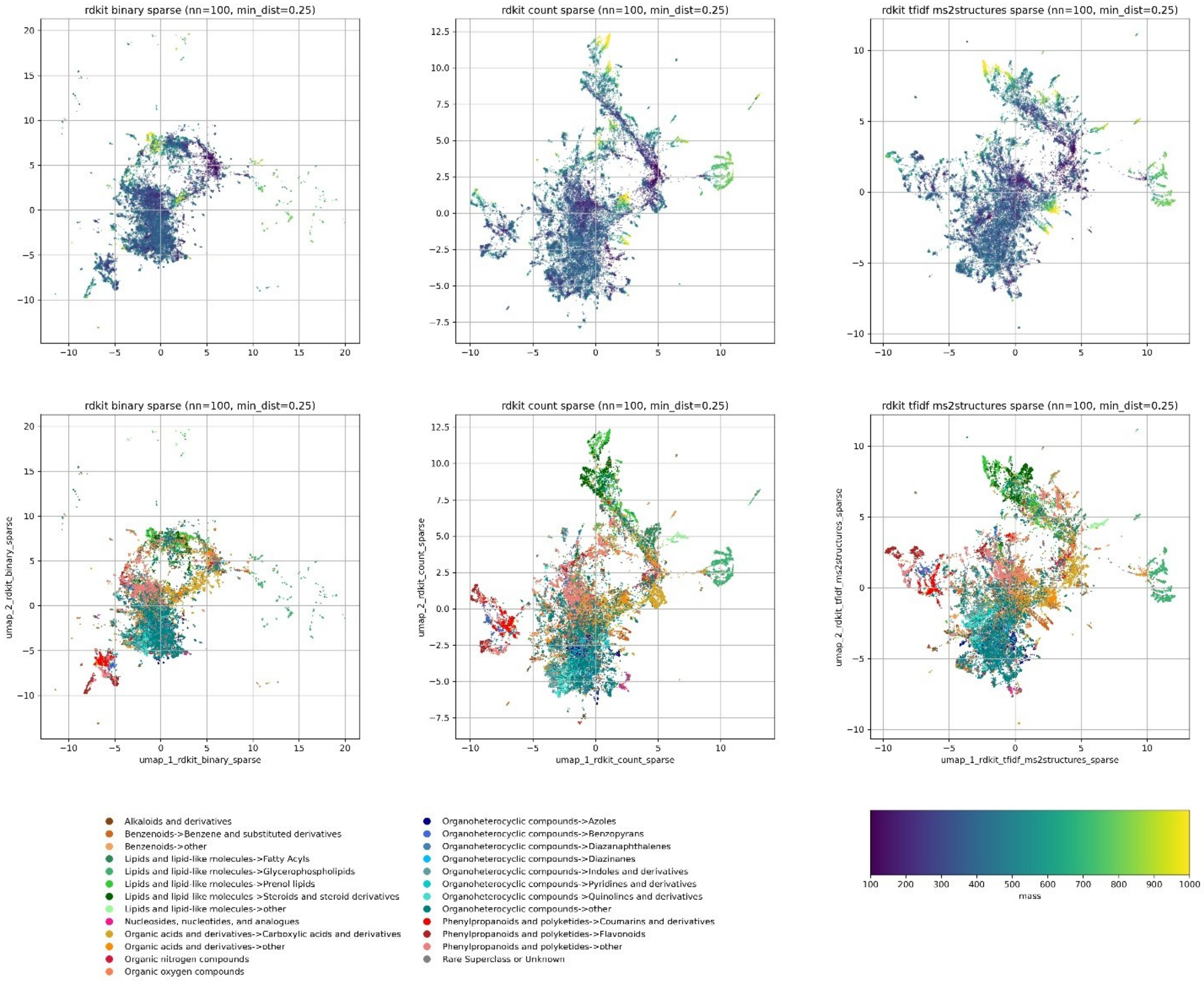


*Figure 12 UMAP-based 2D coordinates of all 38,711 compounds in the ms2structures dataset were used to visualize the chemical space of this dataset according to different similarity measures. UMAP was run using the 100 nearest neighbors (and their similarity scores) computed using Tanimoto scores of RDKit binary sparse fingerprints (left panel), of RDKit count sparse fingerprints (center panel) as well as of IDF-weighted RDKit count sparse fingerprints (right panel). In the top row plots every molecule is colored by its molecular mass (in Da), and in the bottom row plots the molecules are colored based on their chemical class according to Classyfire [42] (less common classes were merged to obtain a limited number of categories for better visualization).*
